# Brain-tuning near criticality in newborns by prenatal experience with language

**DOI:** 10.64898/2026.05.21.726940

**Authors:** Jesus Encinas, Benedetta Mariani, Ramon Guevara, Maria Ortiz-Barajas, Judit Gervain, Samir Suweis, Fabrizio Lombardi

**Affiliations:** Department of Developmental and Social Psychology, University of Padua, Via Venezia, 8, 35131, Padua, Italy; Department of Physics ”Galileo Galilei”, University of Padua, Via F. Marzolo, 8 35131, Padua, Italy; Padova Neuroscience Center, University of Padua,Via G. Orus, 2 35131, Padua, Italy; Integrative Neuroscience and Cognition Center and Centre National de la Recherche Scientifique, Université Paris Cité, Rue des Saint-Pères, 45 75006, Paris, France; Department of Biomedical Sciences, University of Padua,Via U. Bassi 58/B, 35121, Padua, Italy

**Keywords:** brain, speech perception, language development, newborns, criticality, neuronal avalanches, long-range temporal correlations, neuroplasticity

## Abstract

Language development starts early, possibly even in the womb. Recent results suggest that newborns’ neural responses to speech in the prenatally heard language are already different from those to unfamiliar languages. However, the neural dynamics of how these differential responses emerge remains little understood. Here, we hypothesize that they are supported by a functional tuning to criticality—a state that maximizes information transmission, dynamic range, and flexibility—, induced in newborns by the prenatally experienced language. To test this hypothesis, we study resting-state brain activity before and after stimulation with naturally spoken sentences in the prenatal language, French, in a rhythmically similar unfamiliar language, Spanish, as well as in a rhythmically different unfamiliar language, English. We show that the native language elicits brain tuning near criticality, balancing network activity and enhancing temporal correlations. Importantly, we find that the network state is not sensitive to the rhythmically different language, and only partially responds to the rhythmically similar language. These results indicate a stimulus-driven tuning to criticality in newborns, a potential foundation for early neuroplasticity, which relies on prenatal experience and may respond to early functional and developmental needs.

## INTRODUCTION

Criticality describes the emergence of scale-invariant fluctuations in dynamical systems poised near a phase transition, where collective behaviors give rise to scaling laws and universal dynamics across spatial and temporal scales^1^. Criticality has provided a unifying framework for understanding diverse natural phenomena, from earthquakes to cardiac and circadian rhythms^2–6^. Increasing evidence indicates that the human brain likewise operates near a critical state: neuronal ensembles display power-law distributed activity patterns^7–9^, large-scale neuroimaging signals conform to critical dynamics^10–12^, and deviations from criticality have been associated with neurological pathology^13–16^. Given that early neurodevelopment is characterized by profound transitions in neural plasticity accompanying the emergence of perceptual and cognitive functions, it represents a particularly compelling yet largely unexplored context in which to investigate the role of critical dynamics in shaping the brain and its function. Here we test the strong hypothesis that the developing brain’s critical state contributes to young infants’ powerful learning abilities, in particular in the domain of language learning.

Understanding how the human brain supports speech perception and language acquisition from the early moments of life remains a fundamental question in cognitive neuroscience. Newborn infants already possess refined auditory abilities: they can recognize their native language^17,18^, discriminate between unfamiliar languages that are rhythmically different^19–22^, or distinguish words with different stress patterns^23^.

Indeed, language acquisition starts even before birth. Prenatal experience with the language heard in the womb during the last trimester of pregnancy already shapes infants’ speech perception abilities assessed at birth^22,24–28^ In the womb, the speech signal is low-pass filtered by maternal tissues and amniotic fluid^29^. As a result, the high frequencies necessary to identify individual phonemes are suppressed, but prosody, i.e. the rhythm and melody of speech, are preserved and reach the fetus^20,29–31^. This allows fetuses to learn about the prosody of their native language already in utero, which lays the foundations for subsequent language development.

Despite robust evidence for newborns’ sophisticated early speech perception and language learning abilities, the neural mechanisms that support these remain poorly understood. Here, we hypothesize that, already at birth, these mechanisms rely on specific processes at the level of neural dynamics. Specifically, we argue that (i) the brain is already capable of self-tuning its neural population dynamics to “criticality”—a highly sensitive and flexible internal state that maximizes network communication and repertoire for efficient learning and information storage^32–36^—and that (ii) such modulation of the internal state is primarily driven by prenatal experience with language.

To test these hypotheses, we analyze resting-state EEG of prenatally French-exposed newborns (*n* = 33, age: 2.39 days; range 1 to 5 days; 19 girls) before (Silence 1) and after (Silence 2) exposing them to a sequence of naturally spoken sentences in three different languages: French, the prenatal experienced language; Spanish, unfamiliar and rhythmically similar to French; English, unfamiliar and rhythmically different from French. This choice is motivated by the fact that newborns can discriminate between rhythmically different but not between rhythmically similar languages^37,38^, and allows us to assess potential effects of rhythmical similarity. Finding that the newborn brain is in a near-critical state, especially if the brain dynamics are more strongly modulated after stimulation with the native language than with the unfamiliar languages, would indicate that prenatal experience plays a key role in tuning the brain towards criticality, and that criticality may be linked to learning and neuroplasticity.

Criticality is characterized by long-range spatio-temporal correlations and emergent cascades of neural activity with power-law size and duration distributions—indicating absence of characteristic spatial and temporal scales in the dynamics. Such cascades, termed neuronal avalanches^39^, allow for a balanced propagation of neural activity throughout the network, with a branching ratio close to one—i.e., one unit “activates” on average another unit in the network—and sizes and durations that obey specific scaling relationships at criticality^4,40^. Crucially, almost nothing is known about whether the developing brain is close to criticality and, if yes, whether its critical state contributes to young infants’ powerful learning abilities, as may be hypothesized by existing theories on the role of criticality in efficient information transmission^35,41^. In particular, whether newborn infants’ brain responses to language exhibit neuronal avalanches has never been tested before. To address this fundamental unanswered question, our aim has been to identify non-transient, persistent language-dependent modulations of the newborn brain’s internal states. To this effect, we have quantified how close the newborn brain’s resting-state is to criticality by monitoring temporal correlations and evaluating the scaling relationships and branching parameter for neuronal avalanches before and after speech stimulation.

Neuronal avalanches in EEG recordings are defined as continuous sequences of large EEG fluctuations (events) that unfold over multiple electrodes and time bins^42^. We measured avalanche size (*s*) and duration (*T*) distributions, quantified their power-law behavior, and estimated the scaling exponents *τ* and *α*, respectively. We then estimated the exponent *γ* that connects sizes and durations through the relationship ⟨*s*⟩ ~ *T* ^*γ*^, and verified the scaling relation between *τ, α* and *γ*, which is satisfied near criticality^9,42^. To evaluate the effect of speech stimulation on the correlations and internal dynamics of avalanches, we first considered the monotonic relationship between the size of consecutive avalanches as a function of the time elapsed between them, Δ*t*. Near criticality, positive and negative increments in avalanche sizes show balanced, opposite correlations with Δ*t*^43^. Next, we measured the branching parameter, which is defined as the average ratio between the number of events that occur in consecutive time frames and provides an estimate of the number of events “triggered” by a preceding event in the network. This quantity tends to one when the system approaches criticality^44^.

We demonstrate that speech in the prenatally experienced language, but not in the unfamiliar languages, significantly modulates the resting-state dynamics of the newborn brain and drives it closer to criticality. Importantly, we do not analyze brain dynamics *during* stimulation. Existing studies have done that^20,21^. Rather, we investigate how stimulation changes brain dynamics in a lasting manner, even minutes *after* it is over, providing a signature for learning. Specifically, after stimulation with French, all the necessary conditions defining criticality are met: the branching parameter gets significantly closer to one, theoretical scaling relationships for avalanches are satisfied, and temporal correlations are enhanced. These results indicate that in-utero experience with language does not only shape perceptual preferences, but tunes the dynamical regime of the developing brain to maximize flexibility and optimize information processing and learning. This may boost the newborn brain’s ability to process new input from the very first days of extra-uterin life, pointing to an early functional role for criticality in the brain.

## RESULTS

### Neuronal avalanches exhibit properties expected at criticality in newborns exposed to the prenatally experienced language

We analyzed the EEG of 33 newborns recorded during 3 minutes of awake resting-state before and after a sequence of auditory stimuli (Fig. 1a-d; Methods). We refer to the resting-state preceding and following auditory stimulation as Silence 1 (S1) and Silence 2 (S2), respectively. The sequence of auditory stimuli consisted of three separate blocks (Fig. 1c): the language infants heard prenatally (French), a rhythmically similar unfamiliar language (Spanish), and a rhythmically different unfamiliar language (English). The order of the three languages was pseudo-randomized and counterbalanced across infants. Thus roughly one third of the infants heard French as the last language before Silence 2 (henceforth referred to as the French group), one third Spanish (Spanish group) and one third English (English group). To measure neuronal avalanches unfolding across the scalp, we first identified prominent deflections in the continuous signal collected from each sensor (Fig. 1d). We observed that the signal amplitude distributions deviate from a Gaussian fit for values larger than ≈ 2.0 SD in each EEG electrode (Fig. 1e and Fig. S1). This indicates the presence of spatio-temporal correlations, and marks the occurrence of clustered, nearly synchronous neural activity^42,43,45,46^. Thus, for further analysis we considered positive and negative signal deflections that exceeded an amplitude threshold of *θ* = 2 SD (Methods; see SI for robustness of the analysis with respect to *θ* values).

**Figure 1:**
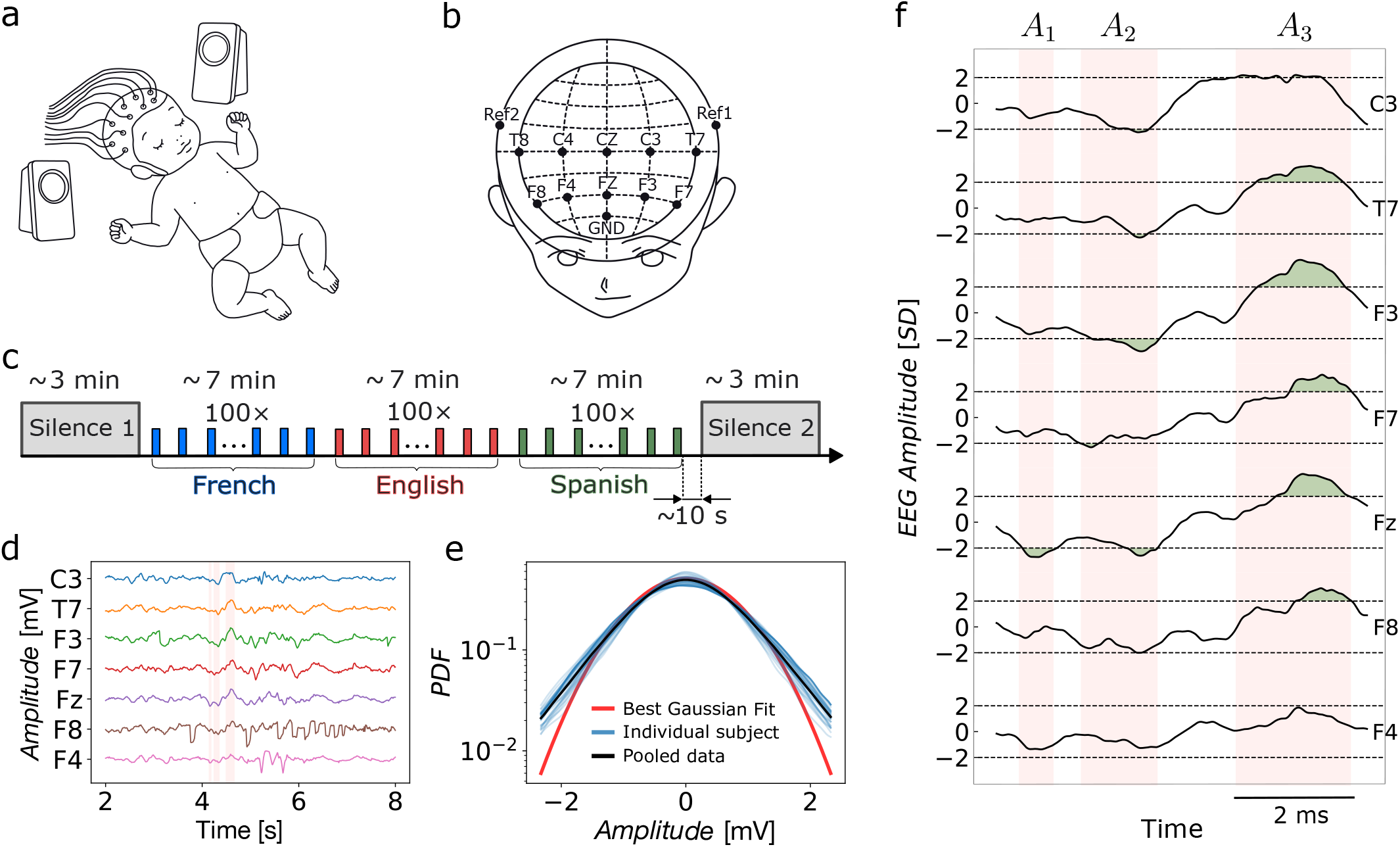
Experimental setup and definition of neuronal avalanches. **a.** Newborns are exposed to auditory stimuli and brain activity is recorded using an EEG cap. **b**. EEG channels layout. 10 channels were used in the study. Channel locations follow the international 10 − 20 system. **c**. Experimental block design. A 3 min silent period (Silence 1) was followed by three blocks of language stimulation (≈ 7 min, 100 stimuli of 2 seconds each), one for each language (French, Spanish, English). After the last language block, resting-state activity was again recorded during a 3 min silent period (Silence 2). **d**. Representative 6 s segments of z-score normalized signal traces for a single subject and 7 selected EEG channels. **e**. Distributions of z-normalized EEG amplitudes for the 33 individual participants in Silence 1 (blue lines), and for data pooled across participants (black line). The red curve represents the best Gaussian fit of the pooled data. The distribution for pooled data starts to deviate from the Gaussian fit at approximately 2 SD. A similar behavior is observed in Silence 2 (Fig. S1). **f**. A neuronal avalanche is defined as a sequence of positive or negative EEG amplitude excursions beyond the threshold *θ* = ± 2SD (green shaded areas) on one or more EEG electrodes. Avalanches are separated by periods with no signal excursions beyond ± 2 SD.

We defined an avalanche as a continuous time interval in which there is at least one excursion beyond threshold in at least one EEG channel (Fig. 1f, red shaded regions). Each avalanche was preceded and followed by time intervals with no other avalanches, i.e. excursions beyond threshold in any EEG channel. We characterized avalanches by their size *s*, and their duration *T*. The size of an avalanche was given by the sum over all channels of the signal absolute values that exceed the threshold *θ*; its duration corresponded to the interval comprised between the first and the last excursions beyond threshold that belonged to the avalanche (Fig. 1f).

We first explored the distributions of avalanche size and duration, *P* (*s*) and *P* (*T*), respectively, during Silence 1. We found that both *P* (*s*) and *P* (*T*) are well described by a power-law on a range of values, i.e. *P* (*s*) ∝ *s*^−*τ*^ and *P* (*T*) ∝ *T*^−*α*^, respectively (Fig. 2). Because small sizes are mostly associated with (local) almost single-electrode events and large sizes are affected by finite size effects^42^(see Fig. S2), we performed the power-law analysis over a limited range of sizes and durations to obtain a reliable estimate of the scaling exponents *τ* and *α*. To robustly identify the onset and offset of the power-law regime, we followed a procedure based on maximum likelihood estimates with false-positive rejection (see Methods and SI, Figs. S3-S5)^47^.

**Figure 2:**
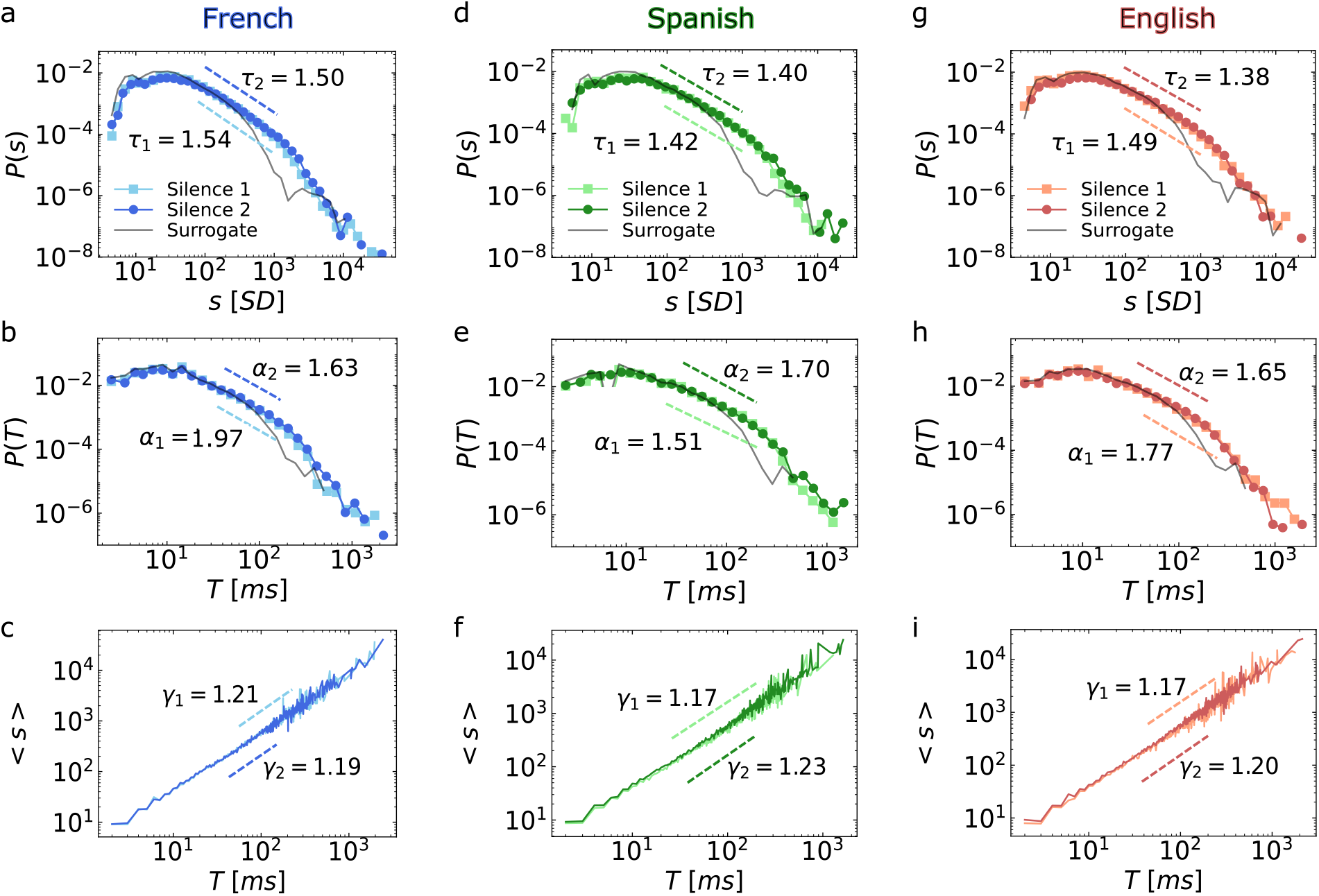
Neuronal avalanches in newborns satisfy predicted scaling relationships after stimulation in the prenatally experienced language. Distributions and scaling relationship for avalanche sizes and durations in Silence 1 and Silence 2. Each column presents, from top to bottom, the distribution of avalanche sizes, *P* (*s*), avalanche durations, *P* (*T*), and the relationship between avalanche sizes and durations, ⟨*s*⟩ ∝ *T*^*γ*^, for the three language groups (pooled data). Avalanche size and duration distributions are consistent with truncated power-laws with exponents *τ* and *α*, respectively. **a-c**. French. Light blue, Silence 1: *τ*_1_ = 1.544 ± 0.007, *α*_1_ = 1.968 ± 0.021, *γ*_1_ = 1.209 ± 0.028 (exponent ± error on the fit); Dark blue, Silence 2: *τ*_2_ = 1.503 ± 0.008, *α*_2_ = 1.627 ± 0.012, *γ*_2_ = 1.186 ± 0.025. **d-f**. Spanish. Light green, Silence 1: *τ*_1_ = 1.416 ± 0.007, *α*_1_ = 1.507 ± 0.009, *γ*_1_ = 1.168 ± 0.017; dark green, Silence 2: *τ*_2_ = 1.397 ± 0.007, *α*_2_ = 1.697 ± 0.014, *γ*_2_ = 1.233 ± 0.016; **g-i**. English. Light red, Silence 1: *τ*_1_ = 1.485 ± 0.008, *α*_1_ = 1.773 ± 0.016, *γ*_1_ = 1.174 ± 0.028; dark red, Silence 2: *τ*_2_ = 1.377 ± 0.006, *α*_2_ = 1.645 + 0.011, *γ*_2_ = 1.204 + 0.019. Black curves: distributions obtained from surrogate data (Methods). Fit in **c**., **f**., and **i**. are linear least-square fit of the form *log* ⟨ *s* ⟩ = *a* · *log T* + *b*. All power-law fits were performed following^47^(Methods).

Our analysis shows that in Silence 1, the exponent *τ* for the size distribution is close to 3/2 in all three groups, while the exponent *α* for the duration distribution ranges between approximately 3 / 2 and 2 (Fig. 2). Comparison of the power-law with an exponential fit by evaluating the log-likelihood ratio 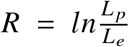 (Methods) between the likelihood *L* _*p*_ for the power law and *L*_*e*_ for the exponential fit indicates that, in the fitted range, the respective power-laws provide a better description of the empirical distributions^48,49^ (Fig. 2 and SI, Table 1 and 2). These results are consistent across subjects (SI, Figs. S6, S7), and stable in a range of *θ* values (Fig. S8): both avalanche size and duration distributions show little variability across participants, and the power-law exponents are consistent with the estimates from pooled distributions (see SI, Table 3). From the analysis of individual participants, we conclude that the variability in *τ* and *α* across groups is not significant and is consistent with the inter-individual variability within groups (SI, Figs. S6, S7 and Table 3)—as expected, since Silence 1 preceded language stimulation and was thus identical across all babies.

The observed power-law exponents are in line with values found in adults in previous studies^42,45,50,51^, indicating that the spatio-temporal dynamics underlying the emergence of neuronal avalanches may already be present immediately after birth. This is further supported by the analysis of surrogate data generated by randomizing the phases of EEG electrode signals (Methods). While preserving the amplitude and autocorrelation of individual signals, this procedure destroys cross-correlations among EEG electrodes. The surrogate analysis showed a significant drop in the probability of large (long) avalanches, which require coordinated electrode activations (i.e., threshold crossing), suggesting that observed avalanche-like behaviors arise from underlying collective dynamics^52^. In this scenario, the average avalanche size is expected to scale as a power-law of the durations, i.e. ⟨*s*⟩ ∝ *T* ^*γ*^, and the exponents *τ, α*, and *γ* to satisfy the relationship^52,53^

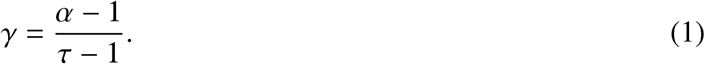

For simplicity, in the following we will refer to the ratio 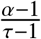 as *R*(*τ, α*). We find that the power-law relation between sizes and durations, ⟨ *s* ⟩ ∝ *T* ^*γ*^, holds in all groups, with *γ*_1_ ≈ 1.2 (Fig. 2 and Fig. S6, S7). However, these *γ* values do not satisfy Eq. 1, as confirmed by the analysis on individual participants (see SI, Table 3). This suggests that, despite hints of scaling in neuronal avalanches, at rest, prior to any stimulation, the newborn brain operates at some distance from criticality. Does language stimulation bring it closer?

To answer this question, we analyzed resting-state activity after stimulation with language (Silence 2, Fig. 2) using the same analysis as for Silence 1. Crucially, the results show that auditory stimulation in the native language, French, modulates avalanche dynamics in such a way that the scaling exponents now satisfy Eq. 1. In particular, language stimulation mainly affects the power-law exponent of avalanche duration distributions, while the exponent of avalanche size distributions remains nearly unchanged in all groups (Fig. 2 and Table 3). In all cases, the analysis of the log-likelihood ratio 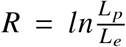 (Methods) indicates that in the fitted range, the power-law provides a better description of the empirical distribution than the exponential fit^48,49^. In the French group, the exponent *α* decreases significantly (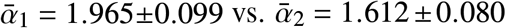 mean ± SEM. *p* = 0.004, n = 12), and Eq. 1 yields 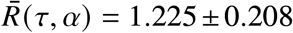 (mean ± SEM), in agreement with *γ*_2_ = 1.210 ± 0.017 (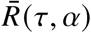 vs. *γ*_2_, *p* = 0.430). Unlike French, Eq. 1 is not satisfied in the Spanish and English groups during Silence 2.

Next, we investigated the temporal shape of avalanches^53^, i.e. the internal temporal structure of avalanches of a given duration (Methods). In short, we averaged the size *s* (*t, T*) at each time *t* (0 < *t* ≤ *T*) elapsed from the beginning of an avalanche across all avalanches of a given duration *T*. The size *s* (*t, T*) corresponds to the number of extreme events within the time interval *t* (Methods). In critical systems, the time course of avalanches follows a universal parabolic shape^53^, namely,

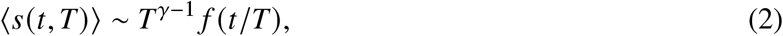

where *f* (*t* / *T*) is an inverted parabola. Equation 2 implies that the avalanche shape is independent of the avalanche duration *T*, which can be verified by plotting the rescaled avalanche time *t* / *T* versus ⟨*s*⟩ *T* ^*γ*−1^ for avalanches of different durations. If Eq. 2 holds, avalanche shapes corresponding to different durations all collapse onto a unique curve, the universal function *f* (*t* /*T*) for a specific *γ*, which is the scaling exponent. In Fig. 3, we show the data collapse for the avalanche shapes in Silence 1 and Silence 2. We observe that avalanches approximate the universal parabolic temporal profile, with scaling exponent *γ* close to those measured in Fig. 2. However, the exponent *γ* that optimizes the shape collapse satisfies the scaling relation in Eq. 1 only in Silence 2 for the French group (Fig. 3).

**Figure 3:**
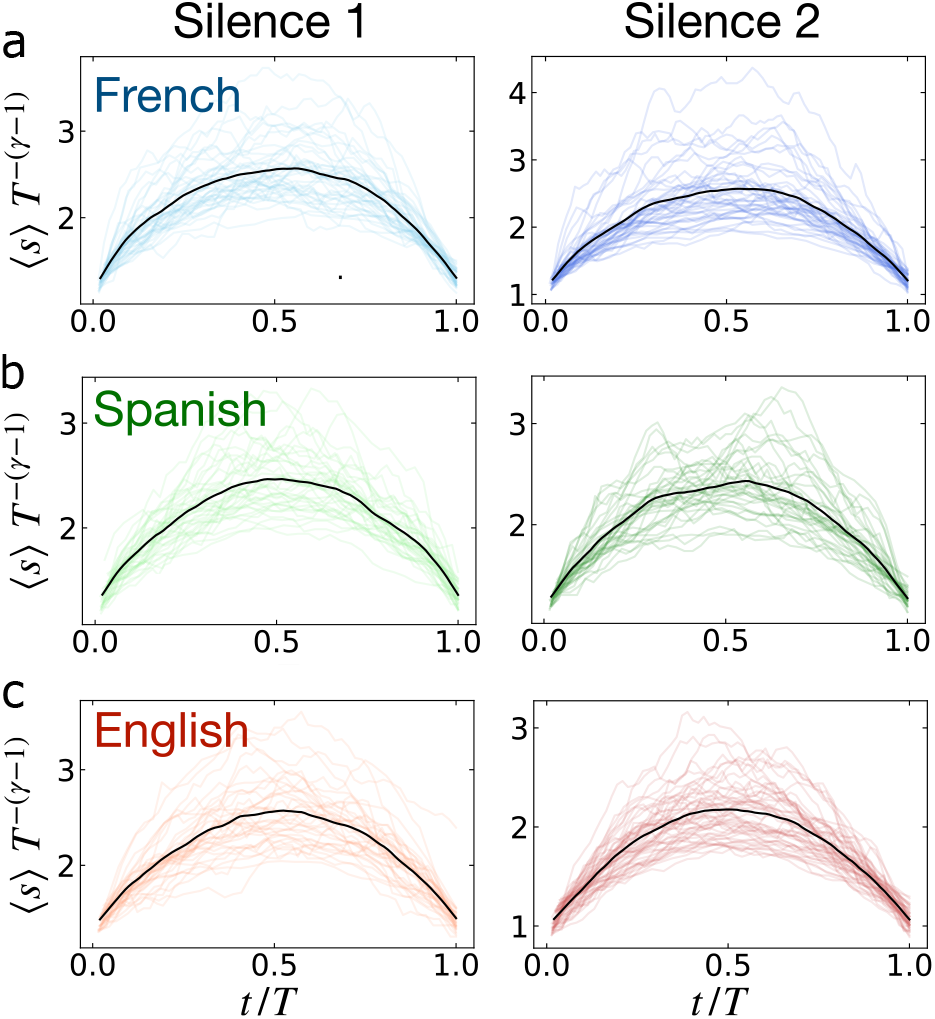
Avalanche shape collapse confirms that relationship between scaling exponents is satisfied only after prenatally experienced language stimulation. Scaling parameters *γ*_1_ and *γ*_2_ for Silence 1 and Silence 2 are estimated through the avalanche shape collapse (Methods). Each colored curve represents the average avalanche profile for a given duration. In each language group and silence type, the average avalanche profile is computed for the avalanche durations within the power-law regime of *P* (*T*) (Fig. 2). The black curve represents the average shape profile over avalanche durations. **a**. French: Silence 1, *γ*_1_ = 1.237±0.018 (fit ± error of the fit); Silence 2, *γ*_2_ = 1.255±0.027. **b**. Spanish: Silence 1, *γ*_1_ = 1.226±0.017; Silence 2, *γ*_2_ = 1.243 ± 0.020. **c**. English: Silence 1, *γ*_1_ = 1.216 ± 0.020; Silence 2, *γ*_2_ = 1.286 ± 0.022.

To characterize the dynamical modulation induced by language stimulation, and thus to further confirm a shift towards criticality, we analyzed the relationship between avalanche size increments {Δ*s*_*i*_} and their corresponding quiet times {Δ*t*_*i*_}. For two consecutive avalanches occurring at time *t*_*i*_ and *t*_*i*+1_, *A*_*i*_ and *A*_*i*+1_, the avalanche size increment is defined as the difference Δ*s*_*i*_ = *s*_*i*+1_ − *s*_*i*_, where *s*_*i*_ is the size of avalanche *A*_*i*_ and *s*_*i*+1_ is the size of the subsequent avalanche *A*_*i*+1_. The quiet time Δ*t*_*i*_ is defined as the difference between the time at which *A*_*i*+1_ starts and the time at which *A*_*i*_ ends. When Δ*s*_*i*_ < 0, avalanche *A*_*i*+1_ is smaller than *A*_*i*_, whereas when Δ*s*_*i*_ > 0 avalanche *A*_*i* + 1_ is larger than *A*_*i*_. We refer to these opposite scenarios as “attenuation” and “amplification”, respectively. Near criticality, Δ*s*_*i*_ and Δ*t*_*i*_ are correlated, and there is a specific balance between attenuation and amplification of neuronal avalanches^43,54,55^.

To assess the attenuation-amplification balance and whether it is modulated by language stimulation, we correlated Δ*s* and Δ*t* using Spearman’s correlation coefficient *ρ* (Δ*s*, Δ*t*), which measures the monotonic relationship between variables (Methods). In Fig. 4, we show the scatter plots between Δ*s*’s and the corresponding Δ*t*’s for the three languages in Silence 1 (Fig. 4a, d, g) and Silence 2 (Fig. 4b, e, h). In the attenuation regime (Δ*s* < 0), we observe that large negative Δ*s*’s tend to occur with short Δ*t*’s, while small negative Δ*s*’s tend to occur with long Δ*t*’s. In this regime, *ρ* (Δ*s* < 0, Δ*t*) is always positive, indicating positive correlations between Δ*s* and Δ*t* (Fig. 4c, f, i). Conversely, in the amplification regime (Δ*s* > 0) large positive Δ*s* tend to couple with short Δ*t*’s, while small positive Δ*s*’s tend to couple with long Δ*t*’s. The Spearman’s correlation coefficient *ρ* (Δ*s* > 0, Δ*t*) is always negative in this regime, indicating significant anti-correlations between positive Δ*s*’s and corresponding Δ*t*’s (Fig. 4c, f, i).

**Figure 4:**
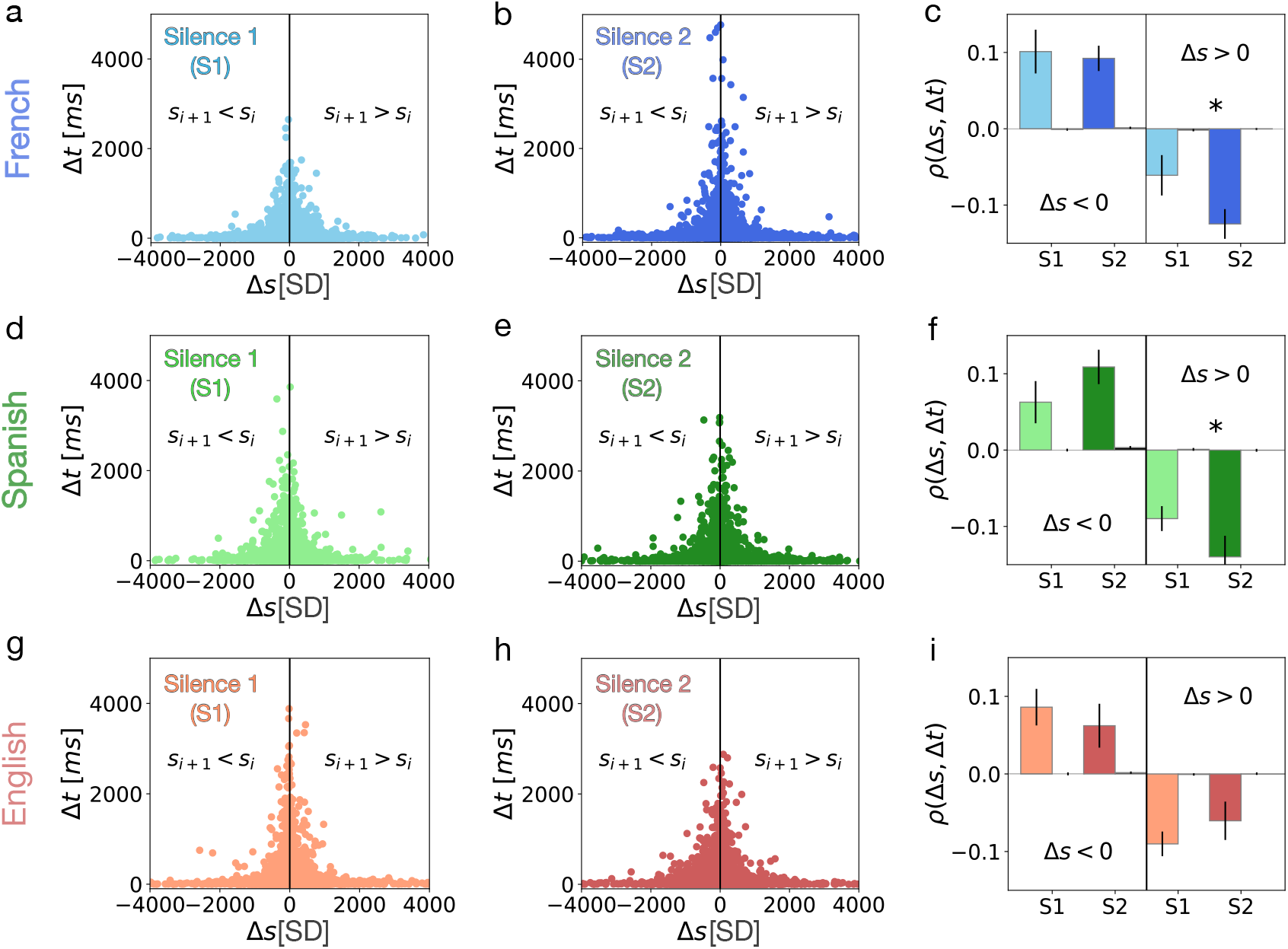
Auditory stimuli in the prenatally experienced language enhance correlations between the sizes of consecutive avalanches. **a-c.** Scatter plot between the size increment, Δ*s* = *s*_*i*+1_ − *s*_*i*_, and the quiet time, 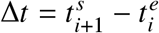, separating two consecutive avalanches *A*_*i*_ and *A*_*i*+1_, for all babies in the French group during Silence 1 (a) and Silence 2 (b). Negative Δ*s*’s are positively correlated with Δ*t*’s, whereas positive Δ*s*’s are anti-correlated with Δ*t*’s (c). Correlations between positive Δ*s* and Δ*t* significantly increase in Silence 2 (*t*-test: *p* = 0.033). **d-f**. A similar behavior is observed for the relationship between Δ*s* and Δ*t* in the Spanish group, with increasing anti-correlations between positive Δ*s*’s and corresponding Δ*t*’s in Silence 2 (Silence 1, *ρ*_1_ (Δ*s* > 0, Δ*t*) = − 0.089 ± 0.016; Silence 2, *ρ*_2(_ Δ*s* > 0, Δ*t*) = −0.139 ± 0.027. *p* = 0.049.). **g-i**. Unlike French and Spanish, English auditory stimuli tend to decrease the strength of correlations between Δ*s*’s and Δ*t*’s, in particular the anti-correlations between positive Δ*s*’s and Δ*t*’s. *ρ*_1_(Δ*s* < 0, Δ*t*) = 0.086 ± 0.023 vs *ρ*_2_(Δ*s* < 0, Δ*t*) = 0.062 ± 0.028; *p* = 0.268. *ρ*_1_(Δ*s* > 0, Δ*t*) = −0.090 ± 0.015 and *ρ*_2_(Δ*s* > 0, Δ*t*) = −0.060 ± 0.024; *p* = 0.207. * = *p* < 0.05.

The comparison between Silence 1 and Silence 2 shows that language stimulation in French enhances correlations between Δ*s* and Δ*t* (Fig. 4a-c). In particular, we observe a significant increase in the amplification regime, Δ*s* > 0, from *ρ*_1_ (Δ*s* > 0, Δ*t*) = − 0.060 ± 0.026 in Silence 1 to *ρ*_2_ (Δ*s* > 0, Δ*t*) = − 0.124 ± 0.019 in Silence 2 (*p* = 0.033), very close to the value observed in adults^43^. This increase may be related to an increase in network sensitivity (excitability), which enhances communication within a network^43^as well as its dynamic range^56^—increasing, for instance, the range of frequencies that can be processed—as expected in states close to criticality. Further, we notice that the increase in correlation strength in the amplification regime compensates for the imbalance between attenuation and amplification in Silence 1 (Fig. 4c)—not observed at criticality^43^. In surrogate datasets obtained by reshuffling avalanche sizes (Methods), the correlations between Δ*s* and Δ*t*, as well as their modulation by language stimulation, are absent (Fig. 4c, f, i, black bars), indicating that the observed correlations arise from and are specific to the underlying avalanche dynamics.

### The prenatally experienced language drives newborn brain dynamics closer to criticality

The relationship between the size of consecutive avalanches suggests that speech in the prenatally experienced language restores the balance between attenuation and amplification of neuronal avalanches. To gain a closer understanding of the modulation induced by language stimulation on resting-state brain dynamics, we next measured how brain activity evolves within avalanches during their propagation over the network. For each avalanche, we recorded the sequence of above-threshold excursions, *n* (*t*), that occurs in each time bin on all electrodes, and estimate the branching parameter *m* (Methods). We refer to each such excursion as an extreme event and to *n* (*t*) as the instantaneous network activity. We note that *n* (*t*) is (approximately) linearly related to avalanche sizes (Fig. S9). A schematic representation of the resulting raster of extreme events from two avalanches is shown in Fig. 4a. The branching parameter is defined as the average of the ratio between the instantaneous network activity at time *t* + 1 and *t* (Methods), and provides a measure of the number of extreme events that, on average, follow an extreme event at a given time *t* (Fig. 4b). The estimate of the branching parameter *m* is indicated as 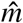^44^. A branching parameter smaller than one indicates that an event “triggers” less than one event on average, and thus the network activity dies out quickly—the smaller the 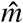, the faster the activity dies out (Fig. 4c). By contrast, 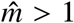 indicates that an event “triggers” more than one event in the following time step, and the activity grows (potentially) unbounded. The 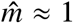 realizes a balanced dynamical regime in which an event “triggers” on average one event in the next time step and the process neither dies out nor grows unbounded (Fig. 4c)—a hallmark of criticality^7,57,58^. In brain dynamics, 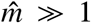 is associated with an excess of network excitation and a sharp increase in very large avalanches^54,58^, whereas 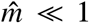 characterizes over-inhibited networks, which exhibit low network activity with reduced sensitivity^59^, a lack of large avalanches, and no scaling in the avalanche size and duration distributions. For neuronal avalanches at criticality, the branching parameter is very close to one, but slightly below, and corresponds to the physiological excitation-inhibition (E/I) balance^42,44,58^.

In Silence 1, we find 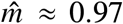 in all groups (Fig. 5d-f). After language stimulation, the branching parameter significantly increases for French and, to a smaller extent, for Spanish, while remains constant for English (Fig. 5d-f). These results hold over a range of threshold values used to define extreme events (Fig. S10). As a baseline of comparison, we calculate the branching parameter on surrogate data generated by randomizing the phase of the EEG signals (Methods; Fig. S11). We obtain a significantly lower value than in the original data in all groups (French: 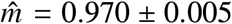 vs 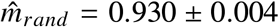, *p* = 10^−7^; Spanish: 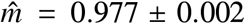 2 vs 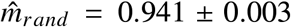, *p* = 10^−7^; English: 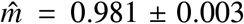 vs 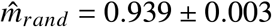, *p* = 10^−7^).

**Figure 5:**
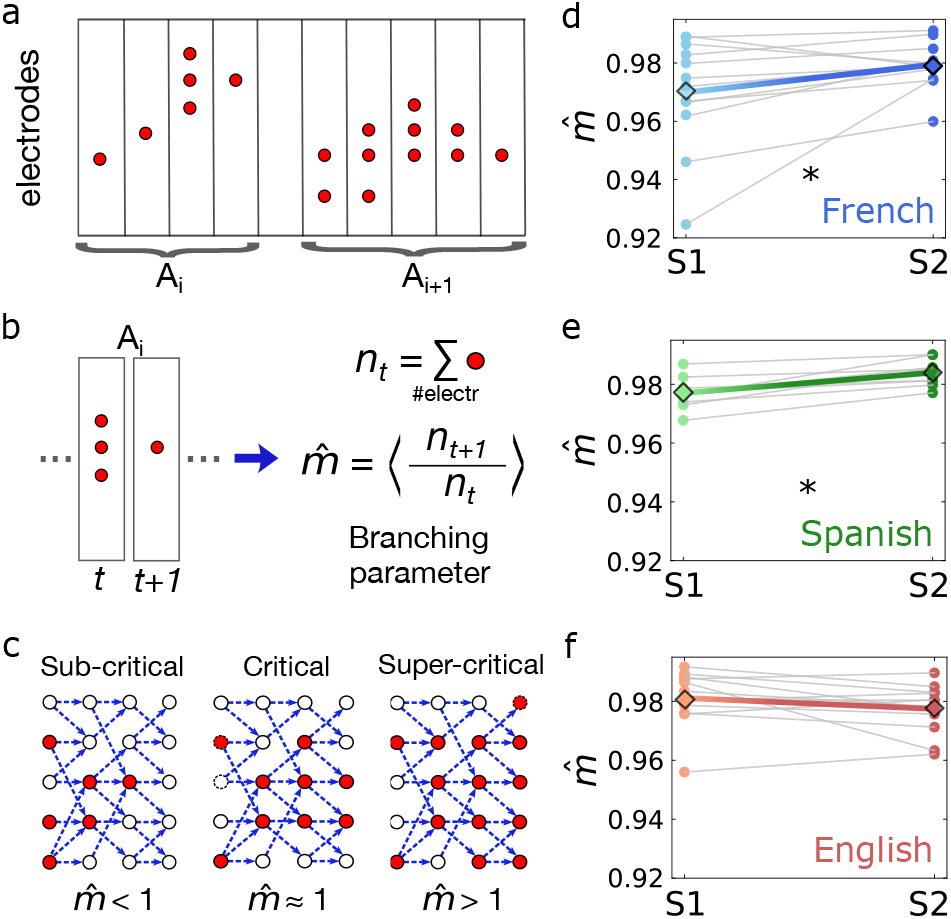
Prenatally experienced language drives the branching parameter of resting-state brain activity closer to the critical value. **a.** Schematic depiction of a sequence of discrete events (red circles) belonging to two consecutive avalanches, *A*_1_ and *A*_2_. Discrete events are defined as positive or negative EEG signal fluctuations that exceed a threshold *θ* (Methods). **b**. In a sequence of events, the branching parameter measure the average number of events triggered at *t* + 1 by an event that has occurred at time *t*, and is formally defined as ⟨*n*_*t*_/*n*_*t*+1_⟩. *n*_*t*_ = number of events measured at the time *t*. **c**. Schematic representation of a sub-critical (left, 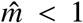), critical (middle, 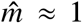) and super-critical (right, *m* > 1) branching process in a network of four layers. In the sub-critical case, the activation rate is smaller that one (one unit activate less than one unit) and the activity dies out quickly. On the contrary, in the supercritical case the activation rate is larger than one (one unit activates multiple units) and network activity grows unbounded (exponentially). Criticality is a balanced regime in which one unit activate (on average) one unit downstream, and the activity can propagate without growing unbounded and potentially causing pathological dynamics. **d-f**. Branching parameter in Silence 1 (S1, before auditory stimuli) and Silence 2 (S2, after auditory stimuli) for individuals in the French (light blue, S1; blue, S2), Spanish (light green, S1; green, S2), and English (light red, S1; red, S2) groups. Auditory stimuli in French cause a significant increase of the branching parameter: from 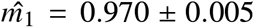 to 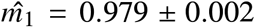 (mean ± SEM; paired *t*-test: *p* = 0.03). A similar behavior, but less pronounced, is observed in the Spanish group, where 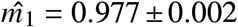 and 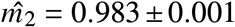 (mean ± SEM; paired *t*-test: *p* = 0.001) in the Spanish group. * = *p* < 0.05. No changes are observed in the English group: 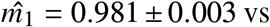. 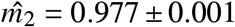 paired *t*-test, *p* > 0.05.

Approaching criticality, we expect the temporal correlation of network activity to decay slower^1^. In Fig. 6, we show the autocorrelation of instantaneous network activity, *C* (*t*), for six representative newborns in the French (four) and Spanish (two) group in Silence 1 and Silence 2. As general features, *C* (*t*) shows an initial exponential decay (*t* < 100 ms), followed by a local peak around ≈ 200 ms—related to theta oscillations^21^—and a slow decay for *t* > 200 ms. However, we observe that in Silence 2 the autocorrelation tends to be stronger, with a much slower decay than in Silence 1 (Fig. 6; see also Figs. S12-S14). Enhanced correlations arise in most participants in the French and Spanish group. We note that the decay of *C* (*t*) is rather slow and much more complex than a simple exponential—a good approximation only at short timescales, *t* ≲ 100 ms—, including multiple timescales and oscillatory trends. For this reason, quantitative comparison between Silence 1 and Silence 2 is rather difficult and may depend on specific model assumptions. In the simple scenario with a fast and slow exponential decay, we did not find significant differences at the group level (Figs. S12-S15). Importantly, we do not observe enhanced correlations in participant in the English group (Fig. S14, S15).

**Figure 6:**
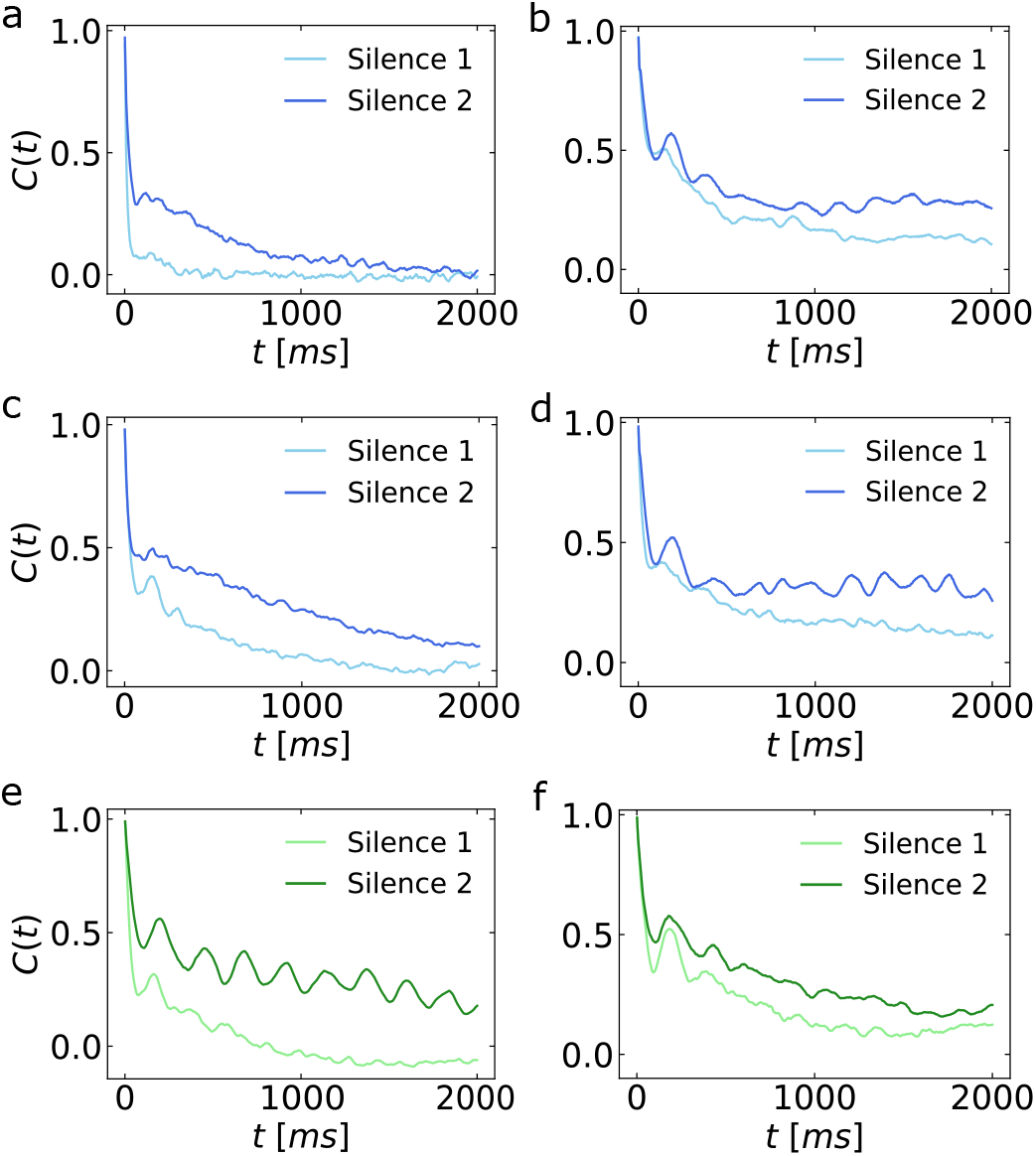
Temporal correlation of collective brain activity increases after stimulation in the prenatally experienced language. a-d. Autocorrelation, *C* (*t*), of the instantaneous network activity for four newborns from the French group in Silence 1 and Silence 2. **e-f**. Autocorrelation, *C* (*t*), of the instantaneous network activity for two newborns from the Spanish group in Silence 1 and Silence 2. The autocorrelation is enhanced during Silence 2, i.e. after auditory stimuli.

### Language-driven modulations of brain dynamics in a neural network model approaching criticality

The analysis of resting-state dynamics provided key evidence in support of our main working hypothesis, namely that external auditory inputs that match the prenatal experience are able to modulate and tune the resting-state brain dynamics to criticality—a state that would support optimal learning^32^. To validate this evidence, here we simulated resting-state activity using a recently proposed neural network model^60^. This model is especially suitable to this end because (i) has a genuine non-equilibrium critical point, (ii) captures local, individual electrode dynamics, including neural oscillations, and (iii) reproduces scale-free neuronal avalanches. Importantly, its analytical tractability offers a direct comparison with EEG data and allows us to infer its two free parameters (Methods).

The model consists of a population of *N* interacting neurons whose dynamics is self-regulated by a time-varying feedback that depends on the ongoing population activity level (Fig. 7a). Neurons are all excitatory, and are simulated as binary units *s*_*i*_ = ±1 (*i* = 1, 2, …, *N, N* = 10^6^in our simulations unless specified differently) that are active when *s*_*i*_ = +1 or inactive when *s*_*i*_ = −1. Here we considered the simplest, fully homogeneous scenario in which neurons interact with each other through synapses of equal strength *J*_*ij*_ = *J* = 1 (see Methods for details). The ongoing network activity is defined as 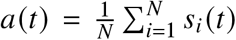, and each neuron experiences a uniform negative feedback *h* that depends on *a* as 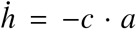, with *c* a positive constant. Neurons *s*_*i*_ are stochastically activated according to the Glauber dynamics, where the new state of neuron *s*_*i*_ is drawn from the marginal Boltzmann-Gibbs distribution 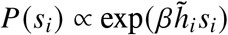, with 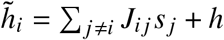, where *β* is a parameter reminiscent of the inverse temperature for an Ising model. For our simulations, we used an all-to-all network connectivity, i.e. *J*_*i j*_ = 1, ∀*i, j* and *i* ≠ *j*. Network behavior is determined by the two parameter *c* and *β*, which control feedback strength and proximity to criticality, respectively. For *c* > 0, the model is driven out of equilibrium and has a critical point at *β* = *β*_*c*_ = 1 ^60,61^. Slightly below the critical point, the model exhibits both oscillations and neuronal avalanches, as we found in newborns.

**Figure 7:**
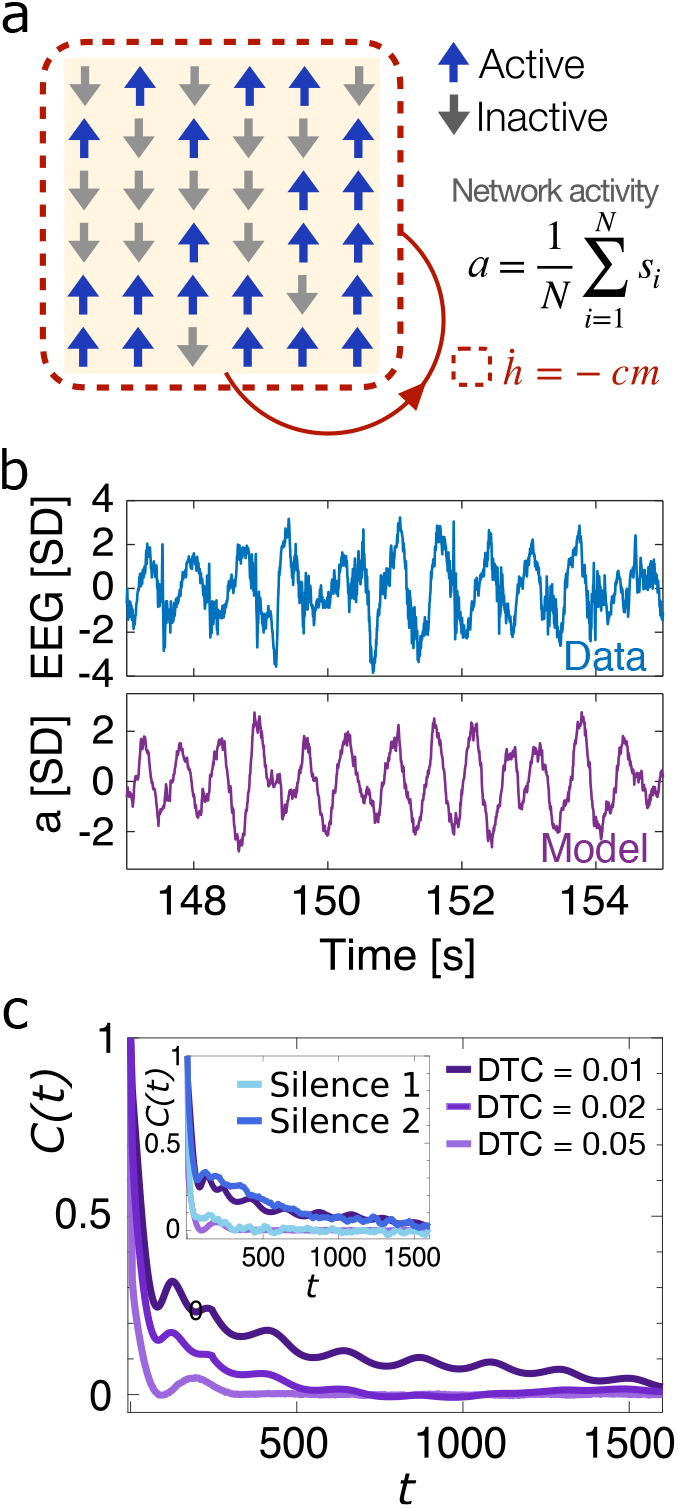
Neural network model of brain dynamics in newborns. **a.** Schematic illustration of the model. Excitatory neurons *s*_*i*_ (*i* = 1, 2, …, *N*) interact with each other through synapses of strength *J*_*i j*_ and experience a time-varying negative feedback *h* (*t*) that mimics inhibition and self-adaptation. Neurons are active when *s*_*i*_ = +1 (up arrows) and inactive when *s*_*i*_ = −1 (down arrows). **b**. Example of an EEG signal (top, blue curve) and the simulated network activity, *a* (*t*), of a model with parameters matched to the data (bottom, violet curve). The model parameters (*β* = 0.99 and *c* = 0.004) are inferred by fitting the analytical form of *C* (*t*) to the autocorrelation estimated from EEG data (SI, Fig. S16). **c**. Autocorrelation function, *C* (*t*), of the model instantaneous network activity, *n* (*t*), for *β* = 0.99, *β* = 0.98 and *β* = 0.95, which correspond to *DTC* = 0.01, 0.02 and 0.05. The increase in correlation strength for *DTC* = 0.01 with respect to *DTC* = 0.02 closely matches the increase observed for several babies in the French group (Fig. 6), suggesting that auditory stimuli in French drive the brain towards criticality. Inset: Model simulations for *DTC* = 0.99 and 0.95 and data from Fig. 6a. Model autocorrelation for *DTC* = 0.01 and 0.02 is smoothed using a moving average filter (100 points) to reduce the oscillatory trend and better visualize the average decay of the correlations.

To estimate the parameters *β* and *c*, we fitted the analytical form of the model autocorrelation to the EEG signal autocorrelation (Methods, Fig. S16). Fig. 7b illustrates the qualitative resemblance between the model and the EEG signal dynamics. Parameters inferred across all sensors and participants strongly concentrate in a narrow range around *β* ≈ 0.99 (French: *β* = 0.9882 ± 0.009), very close to the critical point, and *c* ≈ 0.005 (French: *c* = 0.004 ± 0.002), which we take as baseline values and use for all subsequent data-model comparisons. To make contact with data, we parcel our simulated network into *K* equally-sized disjoint subsystems of *n*_sub_ = *N* / *K* neurons each, and consider each subsystem activity *m*_*μ*_, *μ* = 1, …, *K*, as the equivalent of the EEG signal from a given electrode (Methods). All quantities for the model—network activity, *n* (*t*), avalanche sizes and durations, branching parameter—then follow the same definition as for the data, allowing us to perform direct comparisons as a function of distance to criticality, *DTC* = *β*_*c*_ − *β* (Methods).

In Fig. 7c we show the network activity autocorrelation, *C*_*m*_ (*t*), for the parameter values inferred from empirical data (individual participant, French, Silence 1). The qualitative behavior of *C*_*m*_ (*t*) closely resembles our observation in newborn resting-state activity (Fig. 6): an initial, fast exponential decay, followed by a slower decay. When we decrease *DTC*, the autocorrelation increases—i.e., for a given *t*, correlation is stronger—, and its range becomes progressively longer—as expected when approaching criticality. This behavior closely matches the increase in autocorrelation during Silence 2 in the French group (see Fig. 6), as demonstrated by the comparison of a representative participant in Silence 1 and 2 with simulations at *DTC* = 0.05 and *DTC* = 0.01, respectively (Fig. 7c, inset). We note that as close to criticality as *DTC* = 0.05 (*β* = 0.95) correlations drop to zero at *t* < 100, one order of magnitude faster than for *DTC* = 0.01 (*β* = 0.99) (Fig. 7c). This indicates that small differences in *DTC* have a considerable impact on avalanche dynamics.

Close enough to criticality, we found that neuronal avalanches follow the expected scaling behaviors (Fig. 8a-c). For a *DTC* = 0.01 the distributions of avalanche sizes and durations are well described by power-laws with exponents close to those predicted for the critical branching process (*τ* ≈ 1.5 and *α* ≈ 2), and the mean avalanche size ⟨*s*⟩ scales with durations as a power-law, ⟨*s*⟩ ∝ *T* ^*γ*^. Correspondingly, the shapes of avalanches of different durations collapse onto a unique parabolic shape for *γ* = 1.52, a value close to that obtained from the relationship between ⟨*s*⟩ and *T* (Fig. 8c). Moving away from the critical point, the distributions of avalanche sizes and durations change slowly, showing similar power-law behaviors for 0.01 ≤ *DTC* ≤ 0.05, with 1.5 ≲ *τ* ≲ 2.5 and 2.2 ≲ *α* ≲ 2.8 (Fig. 8a, b). In the range of *DTC* inferred from data, approximately [0.015, 0.01], variability in the scaling behavior of avalanches is limited, as observed in empirical data between Silence 1 and Silence 2. However, for *DTC* ≥ 0.02 we observe a steady increase in the power-law exponents, and both the size and duration distributions approach an exponential behavior for *DTC* ≥ 0.1 (Fig. 8a, b), where the relationship between sizes and durations becomes approximately linear (γ = 1.03 ± 0.01) and the avalanche shape flat (Fig. 8c).

**Figure 8:**
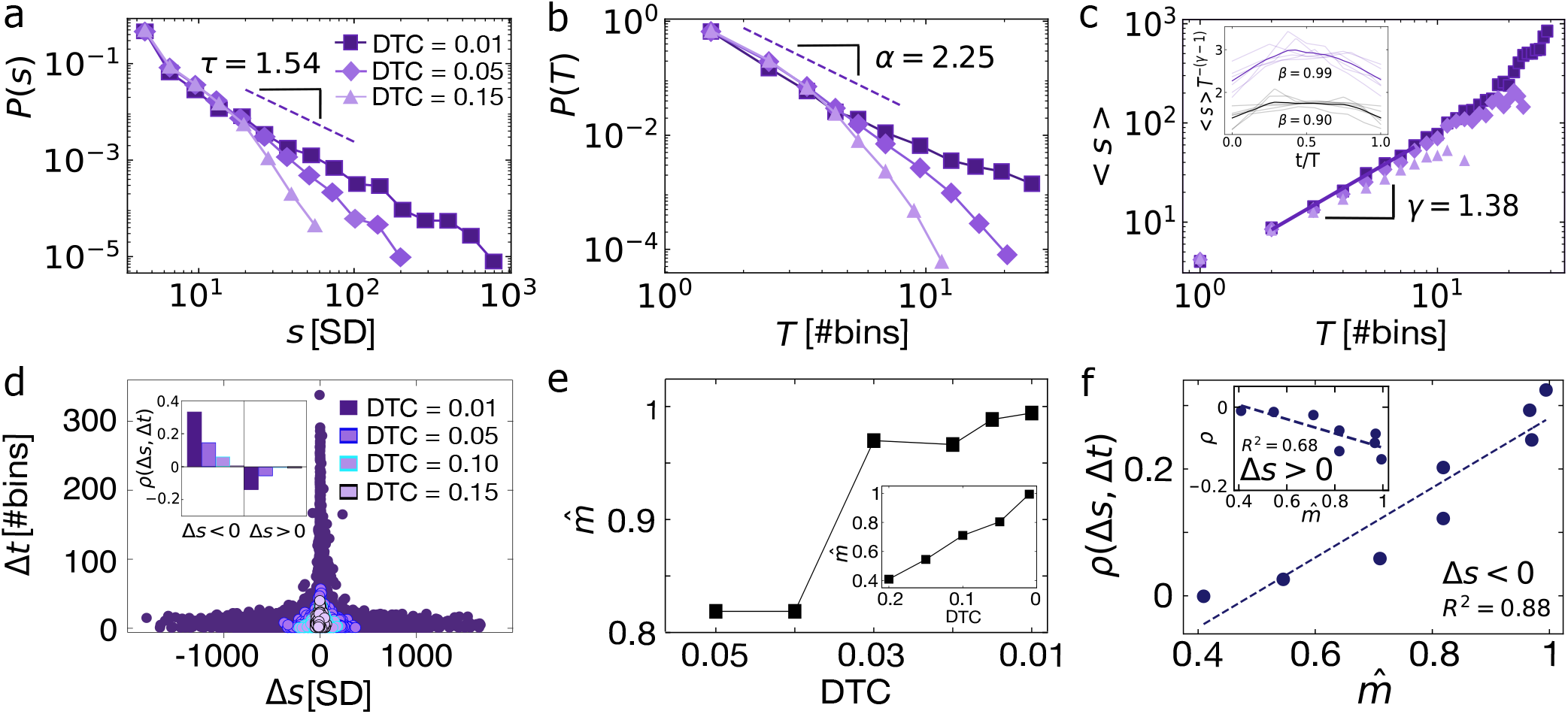
Model simulations indicate that stimulation in prenatally experienced language drives newborns’ resting-state towards criticality. **a.** The distribution of avalanche sizes follows a power-law behavior for the model with 0.01 ≤*DTC* ≤0.05: P (*s*) ∝ *s*^−*τ*^, with *τ* = 1.54 ± 0.01, *τ* = 2.00 ± 0.01, and *τ* = 2.67 ± 0.02 (*R* > 0 and *p* < 10^−5^; see Methods). For *DTC* ≥ 0.05, P (*s*) is better described by an exponential model (*R* < 0, *p* < 10^−5^). **b**. Avalanche duration distributions P (*T*). For 0.01≤ *DTC*≤ 0.05, P (*T*) ∝*T*^−*α*^ with *α* = 2.25 ±0.01, *α* = 2.44± 0.01, and *α* = 2.77± 0.01 (*R* > 0 and *p* < 10^−5^). For *DTC* > 0.05, P (*T*) is better described by an exponential model (*R* < 0, *p* < 10^−5^). **c**. Avalanche sizes scale with their respective durations as ⟨*s* ⟩ ∝*T* ^*γ*^, with a non-trivial exponent *γ* = 1.38± 0.04, *γ* = 1.33± 0.03 and *γ* = 1.30± 0.04 for *DTC* = 0.01, 0.02, 0.05, respectively. For *DTC* 0.1≥ instead, the scaling becomes approximately linear: *γ* = 1, 11± 0.02 for *DTC* = 0.1 and *γ* = 1, 03± 0.01 for *DTC* = 0.15. Purple thick line: linear least square fit of *log s* = *a*· *log T*+ *b*. **d**. Scatter plot between Δ*s* and Δ*t*. For *DTC* = 0.01, the model relationship between Δ*s* and Δ*t* quantitatively resembles the empirical one. When moving away from criticality, the range of Δ*s* and Δ*t* narrows and the correlation between Δ*s* and Δ*t* decreases (inset). **e**. Branching parameter as a function of *DTC*. As *DTC* increases, 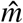 progressively decreases. The branching parameter is close to empirical values 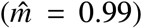 for 0.01 ≤ *DTC* ≤ 0.03, and drops sharply for *DTC* ≥ 0.04. Insets: 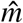 versus *DTC* in the range [0.01, 0.2]. **f**. The correlation strength *ρ*(Δ*s*, Δ*t*), grows with 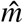. The dashed line is a linear least square fits, *y* = *a* · *x* + *b*, with *a* = 0.55 (*R*^2^ = 0.88, *p* = 0.001). Inset: *ρ*(Δ*s*, Δ*t*) versus 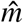 for Δ*s* > 0 (linear fit: *a* = −0.172; *R*^2^ = 0.68, *p* = 0.012). Power-law fits in **a** and **b** were performed following^47,49^. *R* is the ratio between the likelihood of a power-law and an exponential model (Methods).

Turning our attention to avalanche amplification and attenuation dynamics, we found that the relationship between Δ*s* and Δ*t* is also rather sensitive to increases in *DTC*. At *DTC* = 0.01 we observe a very broad range of Δ*s* and Δ*t* and significant correlations between Δ*t* and Δ*s* (Fig. 8d). Both their variability and cross-correlation decrease sharply for *DTC* > 0.02: the range of Δ*s* and Δ*t* narrows considerably (about 4 times smaller at *DTC* = 0.05) and the correlations approach zero at *DTC* = 0.1 (Fig. 8d, inset). The model shows that the variability in Δ*s* and Δ*t*, as well as their reciprocal correlation, are strongly modulated by closeness to criticality.

As avalanche amplification and attenuation are connected to network excitability and balance, we next investigated how the branching parameter, *m*, depends on *DTC*. At the inferred distance to criticality (*DTC* = 0.01), the estimated branching parameter is 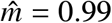, very close to the critical value (*m* = 1) and to the values measured in newborns within the French and Spanish group after stimulation (Fig. 8e). The branching parameter stays close to empirical values for 0.01 ≤ *DTC* ≤ 0.03 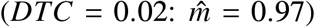, while it drops significantly below empirical values—and even below surrogate values (SI, Fig. S12)—for *DTC* ≤ 0.04, and is approximately 0.5 at *DTC* = 0.15 (Fig. 8e, inset). Importantly, we found that 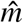 was significantly correlated with *ρ* (Fig. 8f), indicating that the correlation between Δ*s* and Δ*t* is a suitable (and easy to estimate) quantity to track the distance to criticality in empirical data.

## DISCUSSION

Our findings indicate that prenatal experience with language can persistently shape the dynamical regime of the newborn brain. Rather than producing only a transient sensory response during stimulation itself, exposure to the prenatally heard language was followed by resting-state patterns that approached critical dynamics: avalanche statistics became self-consistent, the branching parameter moved closer to 1, and temporal correlations were strengthened. Importantly, this convergence of criticality signatures was not observed after exposure to the rhythmically different unfamiliar language, and only partially for the rhythmically similar unfamiliar language. This evidence argues that resting-state modulation is not explained by auditory stimulation alone, but is linked more specifically to prenatal language experience.

These results are important for two reasons. First, they indicate that neuronal avalanches and scale-free aspects of collective cortical dynamics are already detectable within the first days of life, extending prior work on avalanches beyond adults and older infants^12,42,45,62,63^. Second, they support the idea that closeness to criticality may have functional relevance very early in development. In theoretical and experimental work, critical regimes are associated with enhanced sensitivity, broader dynamic range, and more efficient propagation of activity^33,36,56^. In this context, tuning towards criticality after stimulation in the prenatally heard language may reflect a mechanism by which prenatal experience prepares the newborn brain to process biologically relevant input with greater flexibility. Key in this respect is the observation that the effect appears to be persistent on the timescale of the post-stimulus resting period, not merely locked to the sensory input itself. In the present design, stimulation blocks were separated by silent intervals, and the post-stimulus analysis was performed after an additional delay. Thus, the observed changes in avalanche dynamics and correlation structure occur several minutes after language stimulation, suggesting a shift in the internal state rather than a short-lived evoked response. This interpretation also fits with the broader view that the developing brain does not remain at a fixed operating point, but can be pushed closer to or farther from criticality depending on behavioral or sensory context/needs.

Here, native-language stimulation has been found to bias that operating point selectively. The neural network model strengthens this interpretation by showing that relatively small changes in distance to criticality are sufficient to generate differences in branching ratio, avalanche organization, and temporal correlations comparable to those observed in the data. Importantly, the purpose of the model was not to reproduce exactly every empirical finding, but to show that the direction and magnitude of the observed resting-state modulations are compatible with a network moving within a narrow region near criticality. The model thus provides a mechanistic scaffold for the data rather than a full biological description of the newborn cortex. This framework also offers us insight into the partial effects observed for Spanish. The rhythmically similar unfamiliar language changed some measures, especially the branching parameter, but did not produce the full set of converging signatures seen after French. This intermediate profile is consistent with the idea that rhythmic similarity alone may engage the system to some extent, while prenatal familiarity is required for a more complete and coherent shift in resting-state dynamics. English, by contrast, produced no comparable tuning and in some measures even moved the system away from the pattern expected near criticality. This interpretation meshes well with existing behavioral and brain imaging findings in newborns showing that newborns can clearly distinguish between rhythmically different languages^19^, and show a behavioral preference and increased neural responses to the prenatally heard language^19^, whereas unfamiliar languages rhythmically similar to the native language often produce intermediate results^20–22^.

Despite consistent evidence that the prenatally heard language drives the newborn brain towards criticality, certain limitations need to be acknowledged. EEG recordings were necessarily sparse and short, which is difficult to avoid in developmental neuroimaging studies. These constraints may affect the accuracy of scaling estimates, contributing to the differences between the observed newborn exponents and those reported in adults. In particular, the lower duration exponent could reflect subsampling from the limited number of electrodes, developmental differences in cortical dynamics, or both. Moreover, we could not reliably quantify instantaneous stimulus-induced shifts in proximity to criticality or discriminate language-related differences at that timescale. Future work with denser EEG, larger cohorts, and longitudinal designs will be needed to determine how robust these signatures are across early development, whether they generalize beyond language, and how quickly they emerge or decay. Such studies could also clarify whether the observed differences with adult avalanche statistics mainly reflect genuine developmental differentiation or are mere methodological limitations.

Overall, the present results support the view that prenatal auditory experience does more than just bias perceptual preference: it may transiently set the newborn brain into a dynamical regime better suited for integrating and learning from relevant sensory input. This provides the first evidence that selective tuning toward criticality is already present at the beginning of postnatal life and may represent one mechanism through which early experience shapes cortical function.

## METHODS

### Experimental model and study participant details

Thirty-three newborns aged 1 to 5 days (mean age = 2.55±1.33 days) were selected for the current study from a larger cohort used in a previous study^21^. The experimental protocol was approved by the CER Paris Descartes ethics committee of the Paris Descartes University (now Université Paris Cité). All babies were recruited at the maternity ward of the Robert-Debré Hospital in Paris, and tested during their hospital stay. All parents provided written informed consent and were present during the entire testing session. The inclusion criteria for babies were (i) being full-term and healthy, (ii) having a birth weight > 2.8 kg, (iii) having an Apgar score > 8, (iv) being a maximum of 5 days old, and (v) being born to French native speaker mothers who spoke this language at least 80% of the time during the last three months of pregnancy according to self-report. A total of 54 newborns were tested; 21 participants were excluded from the analysis because of an unfinished test due to fussiness and crying (n=4), technical issues (n=1), poor data quality in one or both silence stages (n=16) (see Pre-processing section for further details). Of the 33 participants analyzed in this study, 16 were girls and 17 were boys. Three groups of babies were formed according to the last language heard in the auditory stimulus stage before the Silence 2 stage: 12 babies heard French, 12 heard English, and 9 heard Spanish.

### Data acquisition

EEG was recorded using an actiCAP-actiCHamp acquisition system from Brain Products GmbH, Gilching, Germany. A 10-channels layout was used, with the lectrodes placed on the following scalp locations: F7, F3, FZ, F4, F8, T7, C3, CZ, C4, and T8. The recording locations were chosen to include the cortical areas where auditory and speech perception-related responses are typically observed in young infants. Channels T7 and T8 were previously labeled as T3 and T4, respectively. Two additional electrodes, one at each mastoid, were used for online reference, and a ground electrode was placed on the forehead. Data were referenced online to the average of the two mastoid channels and were not re-referenced offline. Data were recorded at 500 Hz with an 8 kHz (−3 dB) anti-aliasing filter, and filtered online with a high cutoff filter at 100 Hz, and a low cutoff filter at 0.1 Hz. The electrode impedance was kept below 140 *k*Ω.

### Pre-processing

Data were preprocessed offline using custom Python scripts. Power line noise was removed using a notch filter at 50 Hz, and signals were band-pass filtered between 1 and 100 Hz using a zero-phase-shift Chebyshev filter. Individual filtered EEG signals underwent an artifact rejection process consisting of the following steps. First, channels with amplitudes that exceeded a threshold ± 200*μV* were identified. If this happened in the first or last 30 seconds of one of the 3-minutes resting-state conditions, then only the compromised 30-second data segment was removed. Only channels with a clean recording length of at least 150 seconds were kept. Otherwise, the whole channel was rejected. Second, the average power spectrum (in decibels) of each participant was calculated. If a channel’s power spectrum deviated from the average by more than 4.5 × 10^−5^ dB, that channel was rejected. This procedure was eventually validated by visual inspection to remove any residual artifacts. Subjects with fewer than five valid channels in either silence 1 or silence 2 were excluded from the analysis (n=16). On average, the 33 participants contributed nine clean channels out of the 10 total channels (SD 1.5 for silence 1 and 0.9 for silence 2). All clean channels were low-pass filtered at 50 Hz using a second-order Butterworth filter.

### Extreme events, network activity, and neuronal avalanches

For each subject, EEG signals were normalized to zero mean and unit standard deviation, i.e.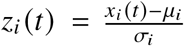, where *x*_*i*_ (*t*) is the clean original signal, *μ*_*i*_ its mean, and *σ*_*i*_ its standard deviation. Positive and negative excursions beyond a threshold *θ* were identified. We refer to them as extreme events or above-threshold excursions. Comparison of the pooled probability distribution of all z-normalized EEG amplitudes with the best Gaussian fit shows that the two distributions start to deviate from each other around ± 2.0SD (Fig. 1c). A Gaussian distribution of amplitudes is expected to be produced from a superposition of uncorrelated sources, and thus is not indicative of relevant extreme events in EEG. For such a reason, one needs to set *θ* ≥ 2SD. Higher values will reduce the number of false positives, but increase the number of false negatives. In this study, the threshold *θ* was set at ± 2.0SD. Extensive robustness analyses were performed to confirm that our key results are stable in a range of *θ* values (see SI, Fig. S8). The network activity, *n* (*t*), is defined as the number of extreme events that occur across all EEG electrodes in a time bin (ϵ = 2 ms). An avalanche is defined as a continuous sequence of time bins in which there is at least an event on any sensor, ending with at least a time bin without events (Fig. 1). The size of an avalanche, *s*, was defined as the sum over all channels of the absolute values of the above-threshold amplitudes^42^.

### Spearman’s correlation coefficient

Given two variables *X* and *Y*, the Spearman’s correlation coefficient is defined as

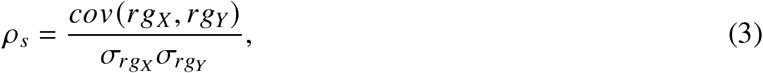

where *rg*_*X*_ and *rg*_*Y*_ are the tied rankings of *X* and *Y*, respectively, 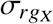 and 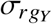 their standard deviations, and *cov* (*rg*_*X*_, *rg*_*Y*_) indicates the covariance between *rg*_*X*_ and *rg*_*Y*_.

### Surrogate signals

Surrogate signals were obtained by random phase shuffling of the original continuous signals. A Fourier transform of each sensor signal was performed, and the corresponding phases were randomized while amplitudes were preserved. The surrogate signals are then obtained by performing an inverse Fourier transform. The random phase shuffling destroys phase synchronization across cortical sites while preserving the linear properties of the original signals, such as power spectral density and two-point correlations^64^. Surrogate signals were used to generate surrogate data for Figs. 2 and S12.

### Surrogate time series for correlations between Δ*s* **and** Δ*t*

To test significance of correlations between consecutive Δ*s* and Δ*t*, a surrogate sequence of avalanche sizes was generated for each subject by randomly reshuffling the original order of avalanche sizes. The Spearman’s correlation coefficient *ρ* (Δ*s*, Δ*t*) between consecutive Δ*s* and Δ*t* was calculated for each surrogate. The average Spearman’s correlation coefficient obtained from all surrogates was then compared with the average correlation coefficient calculated from the original sequences of avalanche sizes and quiet times (Fig. 4).

### Neural network model

The model is composed of a collection of *N* spins *s*_*i*_ = ± 1 (*i* = 1, 2, …, *N*) that interact with each other with a coupling strength 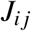. In our analysis, the *N* spins represent excitatory neurons that are active when *s*_*i*_ = +1 or inactive when *s*_*i*_ = −1, and 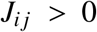. Here, the fully homogeneous scenario is considered, with neurons interacting with each other through synapses of equal strength *J*_*i j*_ = *J* = 1. The *s*_*i*_ are stochastically activated according to the Glauber dynamics, where the state of a neuron is drawn from the marginal Boltzmann-Gibbs distribution

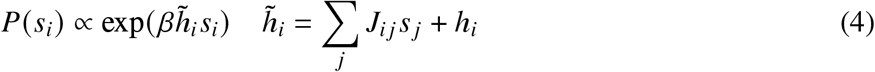

Spins experience an external field *h*, a negative feedback that depends on network activity according to the following equation,

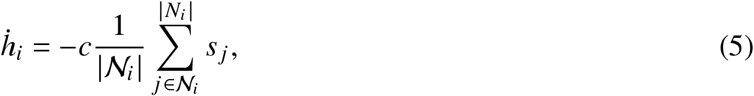

where *c* is a constant that controls the feedback strength, and the sum runs over a neighborhood of the neuron *i* specified by 𝒩_*i*_; index *j* enumerates over all the elements of this neighborhood. In our simulations, the feedback depends on the activity of the entire network^60^.

In the fully-connected case considered here, the model can be solved analytically and a phase diagram can be obtained^60^. In the resonant regime below the critical point (*c* > *c*^*^, *β* < *β*_*c*_), the autocorrelation function, *C*_*a*_ (*τ*), of the ongoing network activity, *a*(*t*), is^60^

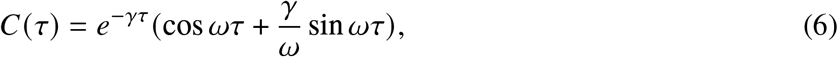

where *γ* = (1 − *β*)/2 is the relaxation time of the system, and 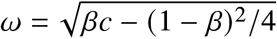 is the characteristic angular frequency of the model.

In our simulations, one time step corresponds to one system sweep — i.e. *N* spin flips — of Monte Carlo updates, and Eq (5) is integrated using Δ*t* = 1/*N*. This choice of timescales for deterministic vs stochastic dynamic is important, as it interpolates between the quasi-equilibrium regime where spins fully equilibrate with respect to the field *h*, and the regime where the field is updated by feedback after each spin-flip and so spins can constantly remain out of equilibrium. Δ*t* is generally much smaller than the characteristic time of the adaptive feedback that is controlled by the parameter *c*.

The parameters *β* and *c* were obtained by minimizing the *L*_2_ error *D* (*c, β*) between the discrete auto-correlation function estimated from data and the analytical expression of the model autocorrelation, i.e.

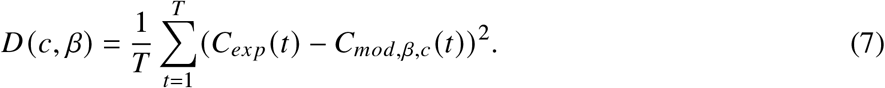

where *C*_*mod,β,c*_ (*t*) is the model autocorrelation with

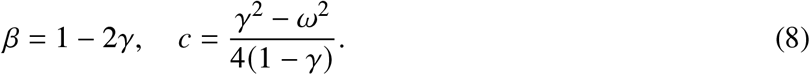

The empirical autocorrelation function, *C*_*ex p*_, was estimated from the z-score normalized electrode signals *z*(*t*):

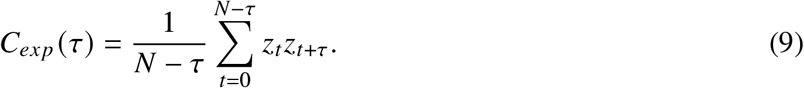

## Quantification and statistical analysis

### Power-law fit

Power law exponents were estimated using a maximum likelihood estimator on double truncated discrete power-laws previously implemented in the NCC Matlab Toolbox^47^. The fitting procedure is briefly summarized here. Consider a power-law distribution

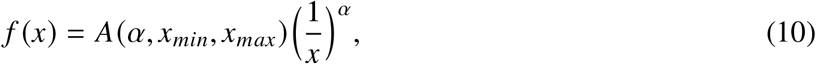

with *x* : [*x* ∈ *Z* : *x*_*min*_ ≤ *x* ≤ *x*_*max*_], *α* the distribution exponent ([*α* ∈ *R* : *α* > 1]), and *A* a normalization constant. The power-law exponent *α* is estimated by maximizing the likelihood function

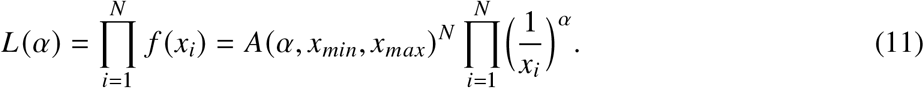

To keep the estimation error below a certain threshold, a recursive search is implemented. The number of recursive steps then coincides with the required decimal precision, e.g, three steps for an error ≥ 10^−3^. First, the log-likelihood 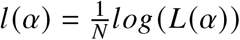 is computed in the interval 1 ≤ *α* ≤ 5 using an increment of 0.1, and the *α* value that maximizes *l α*, (*α* _*f it*.1_), is found. Then, the search continues for *α*_*fit*.1_ − 0.1 ≤ *α* _*fit*.1_ ≤*α*_*fit*.1_ + 0.1 at increments of 0.01 until the value that maximizes *l* (*α* _*fit*.1_) is found. The algorithm continues until it reaches the desired precision. In this study we set the decimal precision to 10^−3^. If at any iteration *α* _*fit*_ is equal to 1 or 5—lower and upper bound of the recursive search—that bound is set to *α* _*fit*_ ^47^.

After an estimate for *α* is found, *N* = 500 synthetic data sets of a power-law distributed variable, *x*_*min*_ < *x* < *x*_*max*_, are generated using a cortical branching model^47^, and the Kolmogorov-Smirnov (KS) statistic is computed for the original data distribution, the distribution obtained from the fitted *α*, and the distribution obtained from the power-law model (PLM). The KS distance between the real data and the fit, (*K S*_*real* −*fit*_), is then compared to the KS distance between the PLM data and the fit, (*K S*_*mod*− *fit*_). If (*K S*_*real* −*fit*_) is at least 10% of the times less than the (*K S*_*mod*− *fit*_) of the PLM (i.e., *p*-value > *p*_*threshold*_ = 0.1, then the truncated power-law is accepted as a good description of the real data^47^.

In our analysis, this procedure was repeated for each language and condition (Silence 1 and Silence 2; pooled data), for a range of *x*_*min*_ and *x*_*max*_ (see SI, Figs. S3-S5). Among the *x*_*min*_ and *x*_*max*_ for which *p* ≥ 0.1, those that minimized the KS distance were selected for the avalanche analysis in Fig. 2. For the selected *x*_*min*_ and *x*_*max*_ in each language and condition, the power law fit was compared to an exponential fit by evaluating the log-likelihood ratio *R* = *lnL* _*p*_/*L*_*e*_, where 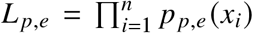 is the likelihood. *R* is positive if the data are more likely to follow a power law distribution, and negative if the data are more likely to follow an exponential distribution. The statistical significance for *R* (*p*-value) was estimated as in^49^.

### Branching parameter

The branching parameter, *m*, was estimated using a spatial subsampling-invariant estimator based on a multiple-regression procedure developed in^44^. A branching process can be approximated with a first-order auto-regressive representation in which the (neural) activity at the next time step, *n* (*t* + 1), depends linearly on the current (neural) activity *n* (*t*), i.e. ⟨ *n* (*t* + 1) | *n* (*t*) ⟩ = *m* · *n* (*t*) + *δ*. In this scenario, the evolution of neural activity *n*(*t*) is governed by *m*, which gives the mean number of ‘neural activations’ triggered by one activation at time *t*. When neural activity is fully sampled, a good estimate of *m* is provided by 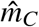, the slope of the linear regression between the time series {*n*(*t*)} and {*n*(*t* + 1)} with *t* = 1, …, *N* − 1, where *N* is the number of time steps. It can be shown that this is not the case when network activity is sub-sampled^44^. To avoid bias in 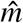, multiple linear regressions of the activity between times *t* and *t* + *k* (*k* = 1, …., *k*_*max*_) are performed, yielding a collection of linear slopes {*r*_*k*_}. The relationship between the activity at time *t* and *t* + *k* is *r*_*k*_ = *b* · *m*^*k*^. The factor *b* is a constant that is generally not known for subsampled activity (b = 1 when the activity is fully sampled) and therefore *m* cannot be obtained directly from any *r*_*k*_ alone. Although subsampling biases *r*_*k*_ by decreasing the mutual dependence between subsequent observations, the temporal decay of *r*_*k*_ is not affected by activity sampling, i.e. *r*_*k*_ ~ *m*^*k*^ = *e*^−*k*Δ*t*/*τ*^ ^44^, where Δ*t* is the time scale of the investigated process (here Δ*t* = 1) and *τ* is the autocorrelation time. Since Δ*t* and *r*_*k*_ are known, 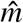 can be obtained from the previous equation for *r*_*k*_, which can be written as *τ* = −Δ*t*/*log*(*m*). After performing *k*_*max*_ linear regression between *n*(*t*) and *n*(*t* + *k*) and obtaining *τ* from an exponential fit of the curve *r*_*k*_ vs. *k*, 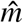 can be easily calculated using the relation between *τ* and *m*. Here, *k*_*max*_ = 150 was used, which roughly corresponds to the cutoff of the initial exponential decay of the autocorrelation of the network activity *n*(*t*).

#### Statistical tests

Pairwise comparisons were conducted using Students two-tailed *t*-test, unless the Shapiro-Wilk test for normality was significant, in which case the non-parametric Mann-Whitney U test was used. A one-way ANOVA test for group comparisons in case data passed the Shapiro-Wilk normality test was used; otherwise, Kruskal-Wallis one-way analysis of variance on ranks (ANOVA on ranks) was used. Multiple pairwise comparisons were performed with the Bonferroni correction. Statistical analyses were performed in Python and MATLAB (Mathworks).

## ACKNOWLEDGMENTS

FL acknowledges support from the European Union’s Horizon research and innovation program under the Marie Sklodowska-Curie Grant Agreement No. 101066790 and from the NextGenerationEU through the grant TAlent in ReSearch@University of Padua – STARS@UNIPD (project BRAINCIP—Brain criticality and information processing). JG acknowledges support from the ERC Consolidator Grant “BabyRhythm 773202”, a FARE grant nr. R204MPRHKE and a PRIN grant nr. 2022WX3FM5 from the Italian Ministry for Universities and Research.

## AUTHOR CONTRIBUTIONS

Conceptualization. J.G., F.L., and S.S.; methodology, J.G., F.L., and S.S.; investigation, J.E., R.G., F.L., B.M., and M.O.-B.; writing-–original draft, J.E., J.G., F.L., and S.S.; writing-–review & editing, all authors; funding acquisition, J.G., F.L. and S.S.; supervision, J.G., F.L., and S.S.

## DECLARATION OF INTERESTS

The authors declare no competing interests.

## DATA AND CODE AVAILABILITY

Data reported in this paper will be shared upon request. This study does not report original code or original data. Additional information required to reanalyze the data reported in this document is available upon request.

## SUPPLEMENTAL INFORMATION INDEX

Figures S1-S16 and their legends.

Table 1.

Table 2.

Table 3.

## Notes

### Competing Interest Statement

The authors have declared no competing interest.

