## Supplemental figures and tables for "Brain-tuning near criticality in newborns by prenatal experience with language"

#### Supplementary figures

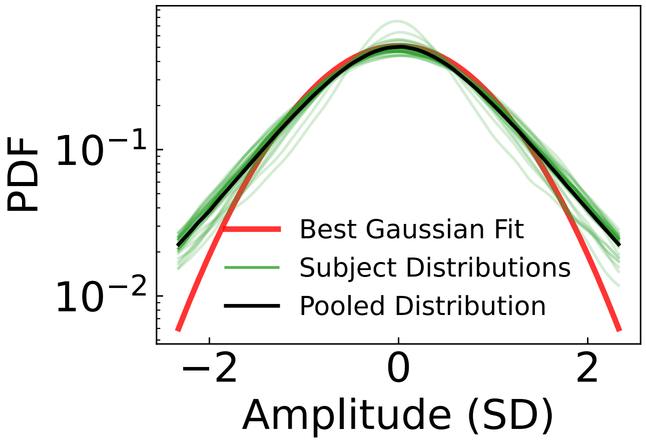


Fig. S1: **Distributions of z-normalized EEG amplitudes in Silence 2.** The probability density distributions of the 33 individual subjects are presented in light green. The black curve is the amplitude distribution for the pooled subjects. The red curve represents the best Gaussian fit on the pooled data. The distribution for pooled data begins to deviate from the Gaussian fit at approximately 2 standard deviations (SD).

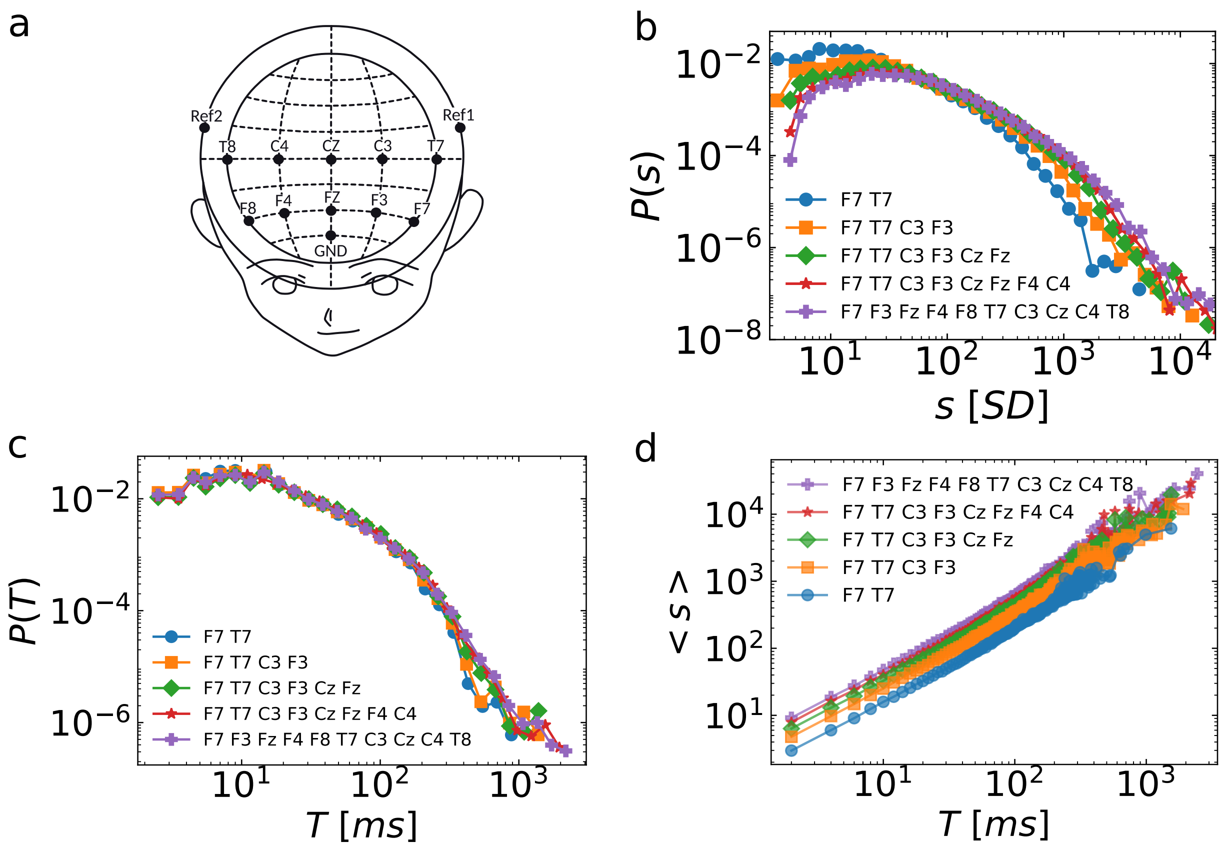


Fig. S2: **Finite size effects on the avalanche size and durations distributions and on the relation** $< s > \propto T^{\gamma}$. **a** EEG channels layout. **b** Removing electrodes from the analysis (systematically, two at a time, from the right to the left hemisphere) the maximum size and the cutoff of the scaling regime decrease, while the probability of smaller avalanche sizes increases. **c** Removing electrodes from the analysis has little influence of the avalanche duration distribution($P(T)$). This may be due to the fact that periods with above threhold excursions have a similar durations across sensors, signaling a fully cooperative behavior. **c** Average size as a function of a given duration, $T$, under systematic channel removal from right to the left hemisphere. (violet plus) 10 channels, (red star) 8 channels, (green diamond) 6 channels, (orange square) 4 channels, and (blue circle) 2 channels. Similar results were obtained when starting from the left to the right hemispheres.

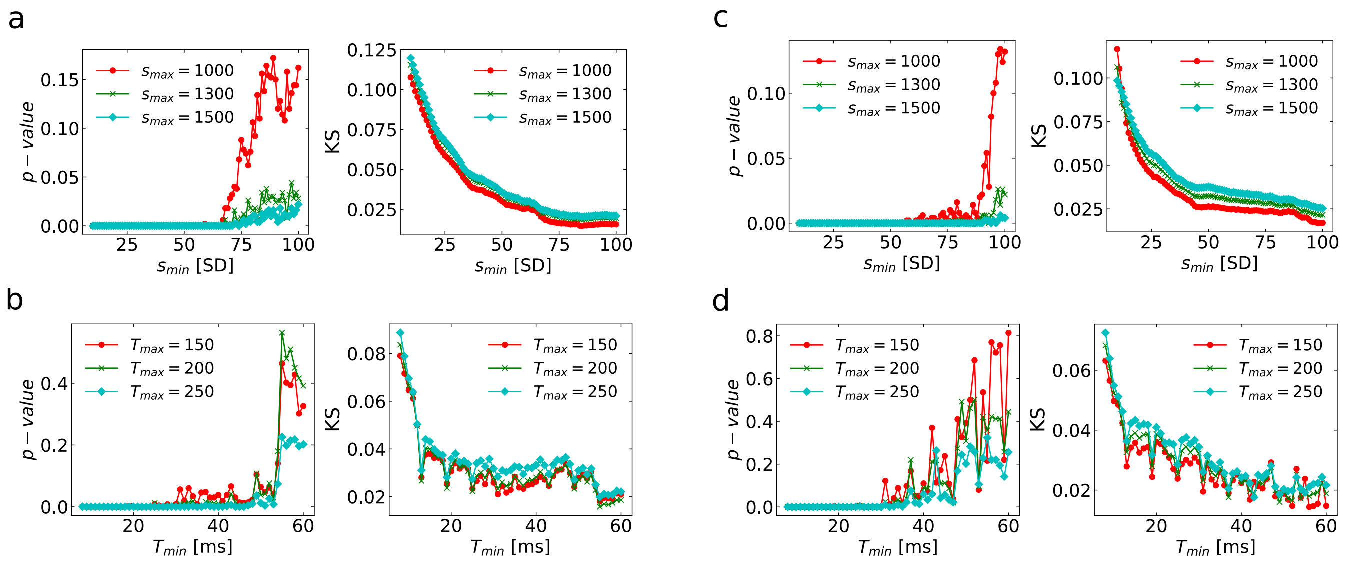


Fig. S3: **Estimation of power law range for avalanche size and duration distributions in the French group. a.-b.** Estimation in Silence 1 of the lower and upper avalanche sizes (durations) $s_{min}(T_{min})$ and $s_{max}(T_{max})$, respectively, for which the empirical avalanche size (duration) distribution is best fitted by a power-law. The $p$-value vs. $s_{min}(T_{min})$ figures show the probability that the Kolmogorov-Smirnov statistic (KS) between the empirical dataset and its double truncated power-law fit is lower than the KS between a power-law model and its fit over 500 realizations. The KS vs. $s_{min}(T_{min})$ plots show the KS value between the empirical dataset and its double truncated power-law fit for several truncating limits in avalanche sizes (durations). Pairs of truncation limits have the minimum KS value among those for which $p - value > 0.1$. In both silences, $s_{min}$ varies in the range $1 \leq s_{min} \leq 100$ and $s_{max} = 1000$ (red circles), 1300 (green $\times$), and 1500 (cyan diamonds). For avalanche durations, $2 \leq T_{min} \leq 60$ ms and $T_{max} = 150$ (red circles), 200 (green $\times$), and 250 ms (cyan diamonds). **a** Silence 1 power-law range estimates for size distribution: $s_{min} = 80$ SD and $s_{max} = 1000$ SD, with $p - value = 0.106$ and $KS = 0.015$. **b** Silence 1 power-law range estimates for duration distribution: $T_{min} = 55$ ms and $T_{max} = 200$ ms, with $p - value = 0.564$ and $KS = 0.015$. **c** Silence 2 power-law range estimates for size distribution: $s_{min} = 100$ SD and $s_{max} = 1000$ SD, with $p - value = 0.114$ and $KS = 0.017$. **d** Silence 2 power-law range estimates for duration distribution: $T_{min} = 42$ ms, $T_{max} = 150$ ms , with $p - value = 0.37$ and $KS = 0.016$.

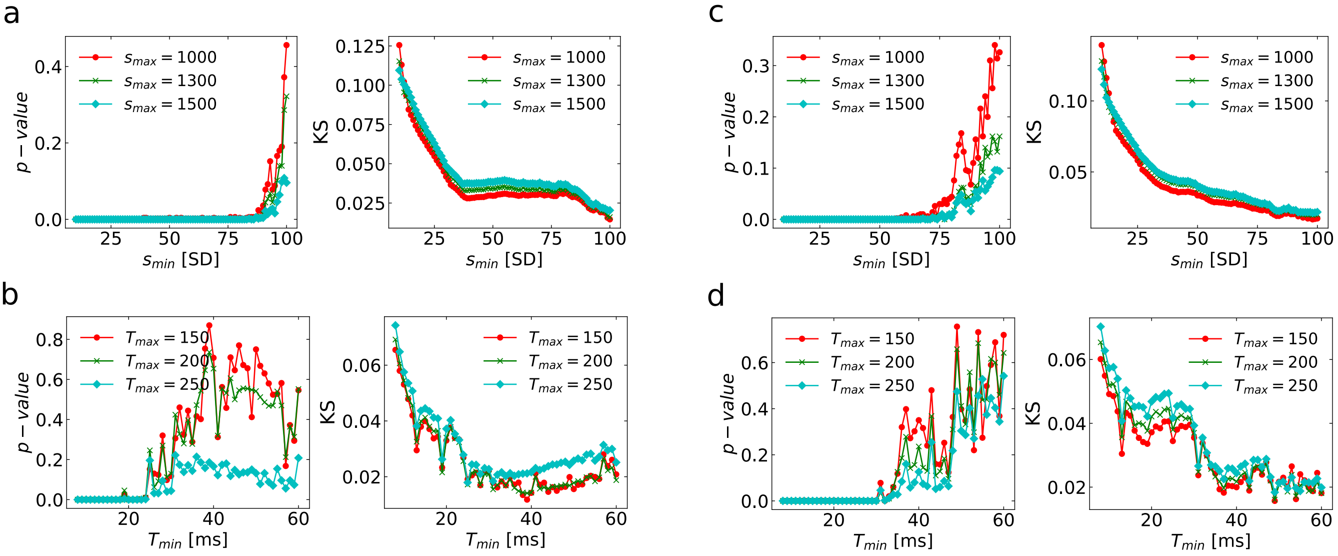


Fig. S4: **Estimation of power-law range for avalanche size and duration distributions for the Spanish group.** Estimation follows the steps described in Fig. S3 and STAR Methods. **a** Silence 1 power-law range estimates for size distribution: $s_{min} = 93$ SD and $s_{max} = 1000$ SD, with $p-value = 0.152$ and $KS = 0.020$. **b** Silence 1 power-law range estimates for duration distribution: $T_{min} = 25$ ms and $T_{max} = 200$ ms, with $p-value = 0.246$ and $KS = 0.017$. **c** Silence 2 power-law range estimates for size distribution: $s_{min} = 82$ SD and $s_{max} = 1000$ SD, with $p-value = 0.124$ and $KS = 0.019$. **d** Silence 2 power-law range estimates for duration distribution: $T_{min} = 37$ ms and $T_{max} = 200$ ms , with $p-value = 0.278$ and $KS = 0.019$.

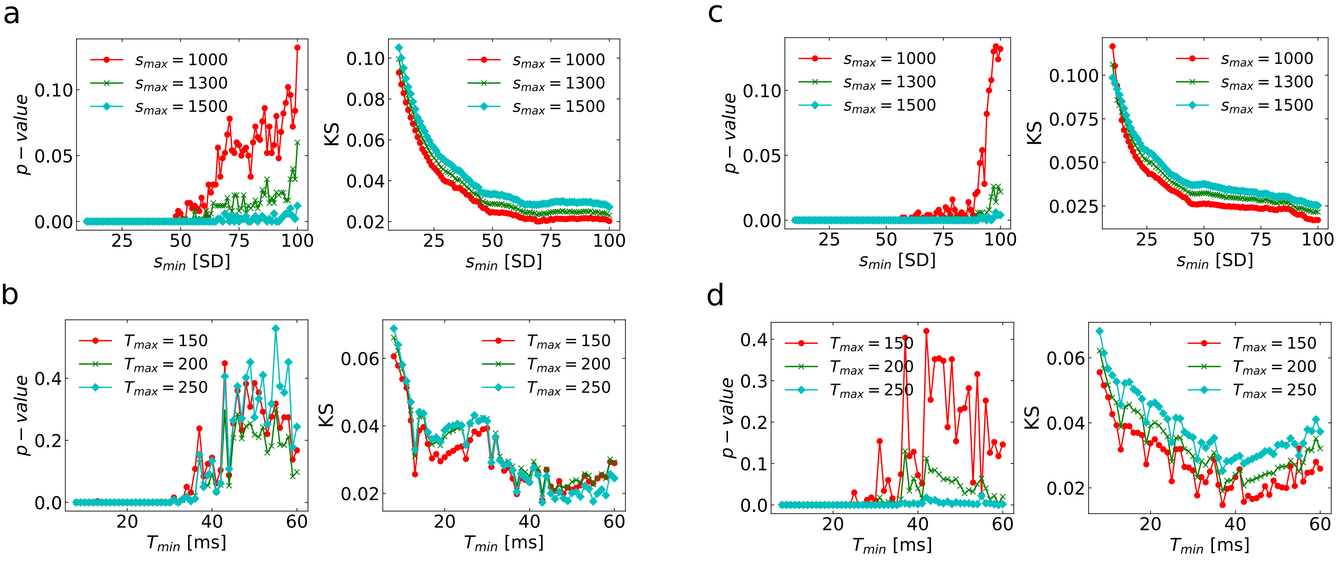


Fig. S5: **Estimation of power-law range for avalanche size and duration distributions for the English group.** Estimation follows the steps described in Fig. S3 and STAR Methods. **a** Silence 1 power-law range estimates for size distribution: $s_{min} = 93$ SD and $s_{max} = 1000$ SD, with $p-value = 0.102$ and $KS = 0.021$. **b** Silence 1 power-law range estimates for duration distribution: $T_{min} = 43$ ms and $T_{max} = 250$ ms, with $p-value = 0.406$ and $KS = 0.017$. **c** Silence 2 power-law range estimates for size distribution $s_{min} = 95$ SD and $s_{max} = 1000$ SD, with $p-value = 0.1$ and $KS = 0.017$. **d** Silence 2 power-law range estimates for duration distribution: $T_{min} = 37$ ms, $T_{max} = 200$ ms , with $p-value = 0.130$ and $KS = 0.018$.

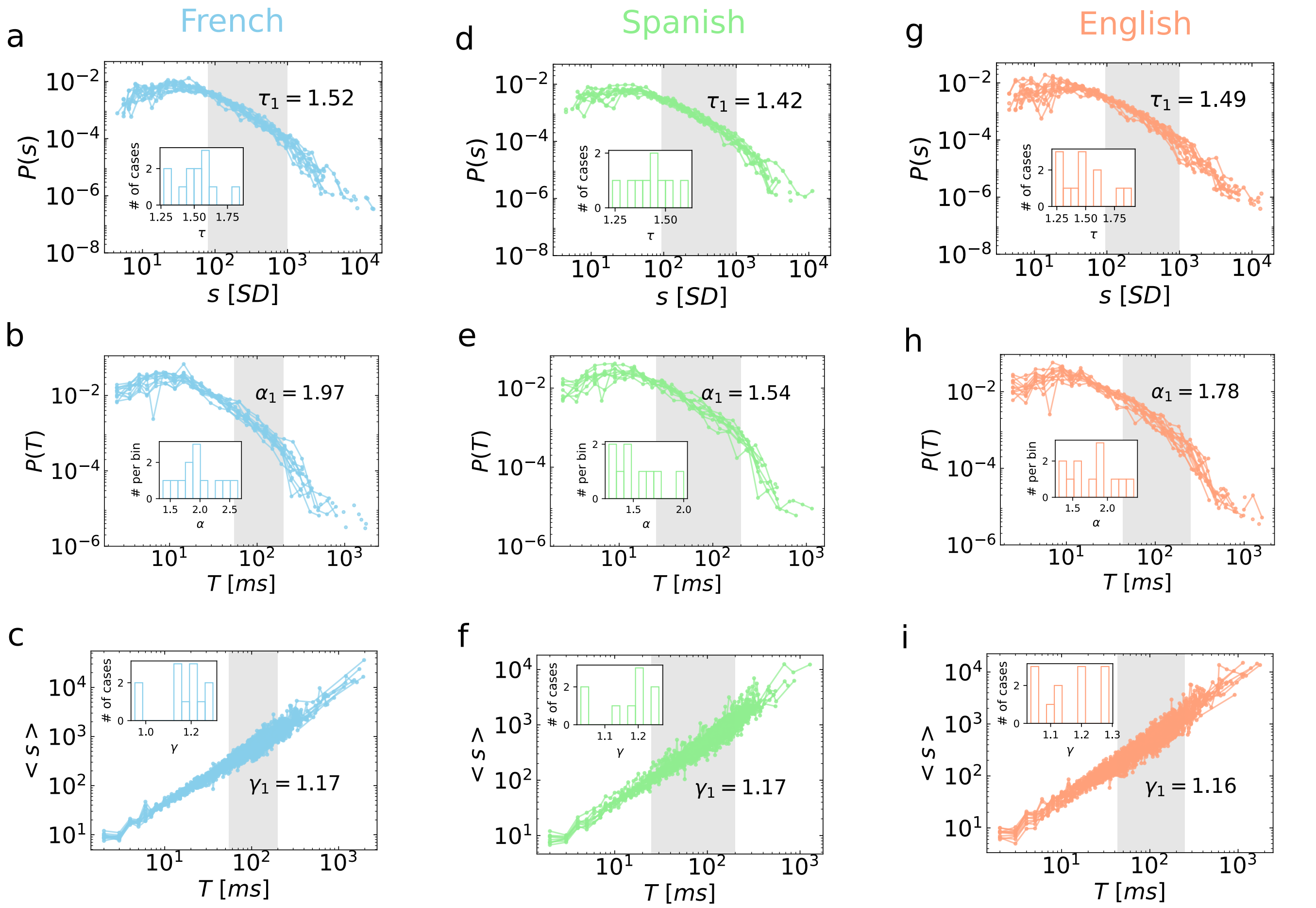


Fig. S6: **Distributions and scaling relationship for avalanche sizes and durations in silence 1 of each language across subjects.** From top to bottom, distributions of avalanche sizes $P(s)$, avalanche durations, $P(T)$, and the relationship between avalanche sizes and durations, $\langle s \rangle \propto T^{\gamma}$, for the three different languages analyzed (individual subjects). Avalanche size and duration distributions are consistent with truncated power laws in the range highlighted in gray, with exponents $\tau$ and $\alpha$, respectively. **a-c.** French, 12 subjects: $\tau_1 = 1.522 \pm 0.043$, $\alpha_1 = 1.965 \pm 0.100$, and $\gamma_1 = 1.165 \pm 0.029$ (mean ± SEM); **d-f.** Spanish, 9 subjects: $\tau_1 = 1.425 \pm 0.034$, $\alpha_1 = 1.536 \pm 0.075$, and $\gamma_1 = 1.166 \pm 0.026$ (mean ± SEM). **g-i.** English, 12 subjects: $\tau_1 = 1.493 \pm 0.055$, $\alpha_1 = 1.781 \pm 0.095$, and $\gamma_1 = 1.162 \pm 0.026$ (mean ± SEM).

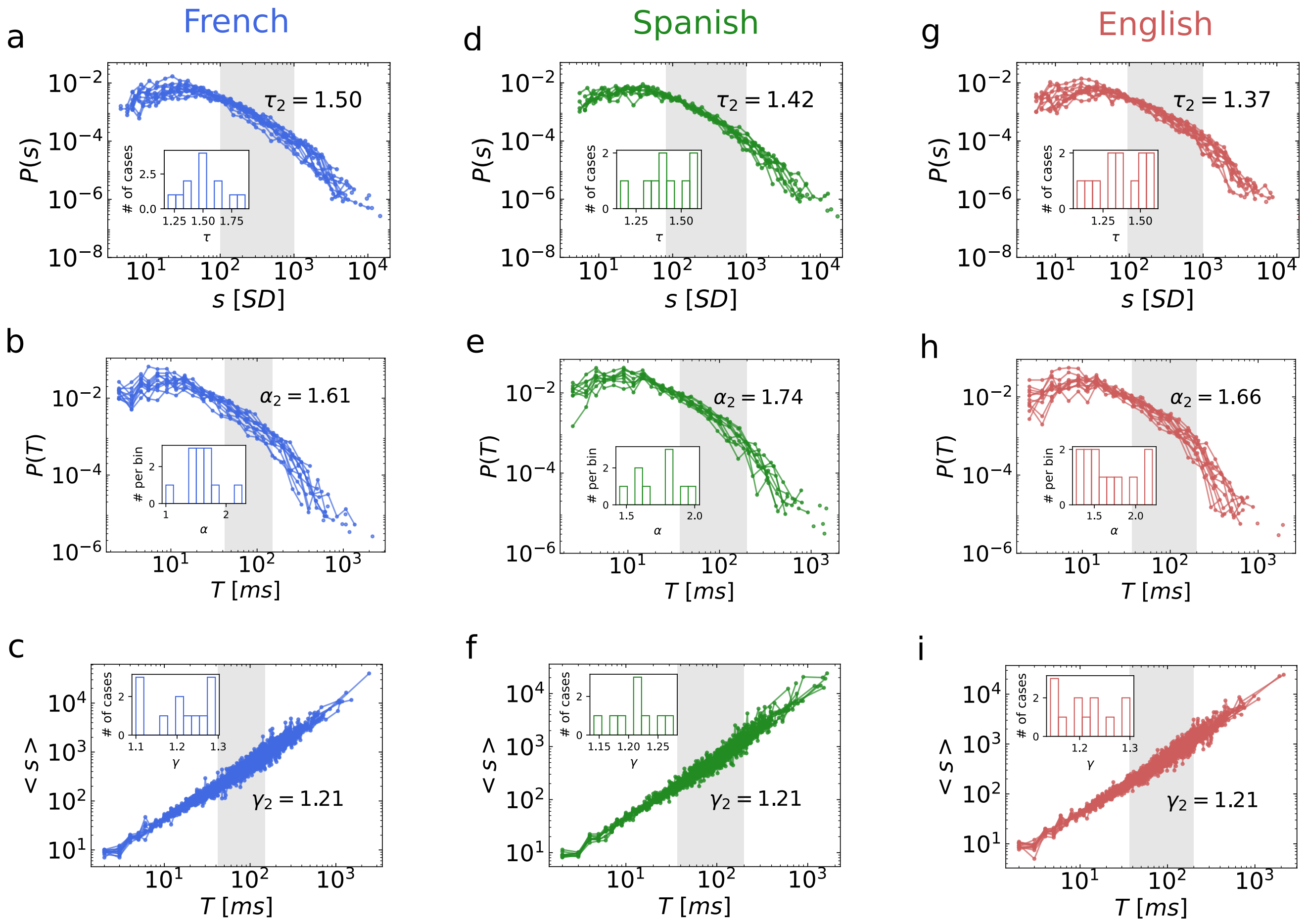


Fig. S7: **Distributions and scaling relationship for avalanche sizes and durations in silence 2 of each language across subjects**. From top to bottom, distributions of avalanche sizes $P(s)$, avalanche durations, $P(T)$, and the relationship between avalanche sizes and durations, $\langle s \rangle \propto T^{\gamma}$, for the three different languages analyzed (individual subject data). Avalanche size and duration distributions are consistent with truncated power laws in the range highlighted in gray, with exponents $\tau$ and $\alpha$, respectively. **a-c.** French, 12 subjects: $\tau_2 = 1.500 \pm 0.054$, $\alpha_2 = 1.613 \pm 0.80$, and $\gamma_2 = 1.207 \pm 0.019$ (main ± SEM); **d-f.** Spanish, 9 subjects: $\tau_2 = 1.417 \pm 0.042$, $\alpha_2 = 1.742 \pm 0.059$, and $\gamma_2 = 1.212 \pm 0.013$ (main ± SEM). **g-i.** English, 12 subjects: $\tau_2 = 1.370 \pm 0.047$, $\alpha_2 = 1.658 \pm 0.087$, and $\gamma_2 = 1.211 \pm 0.015$ (main ± SEM).

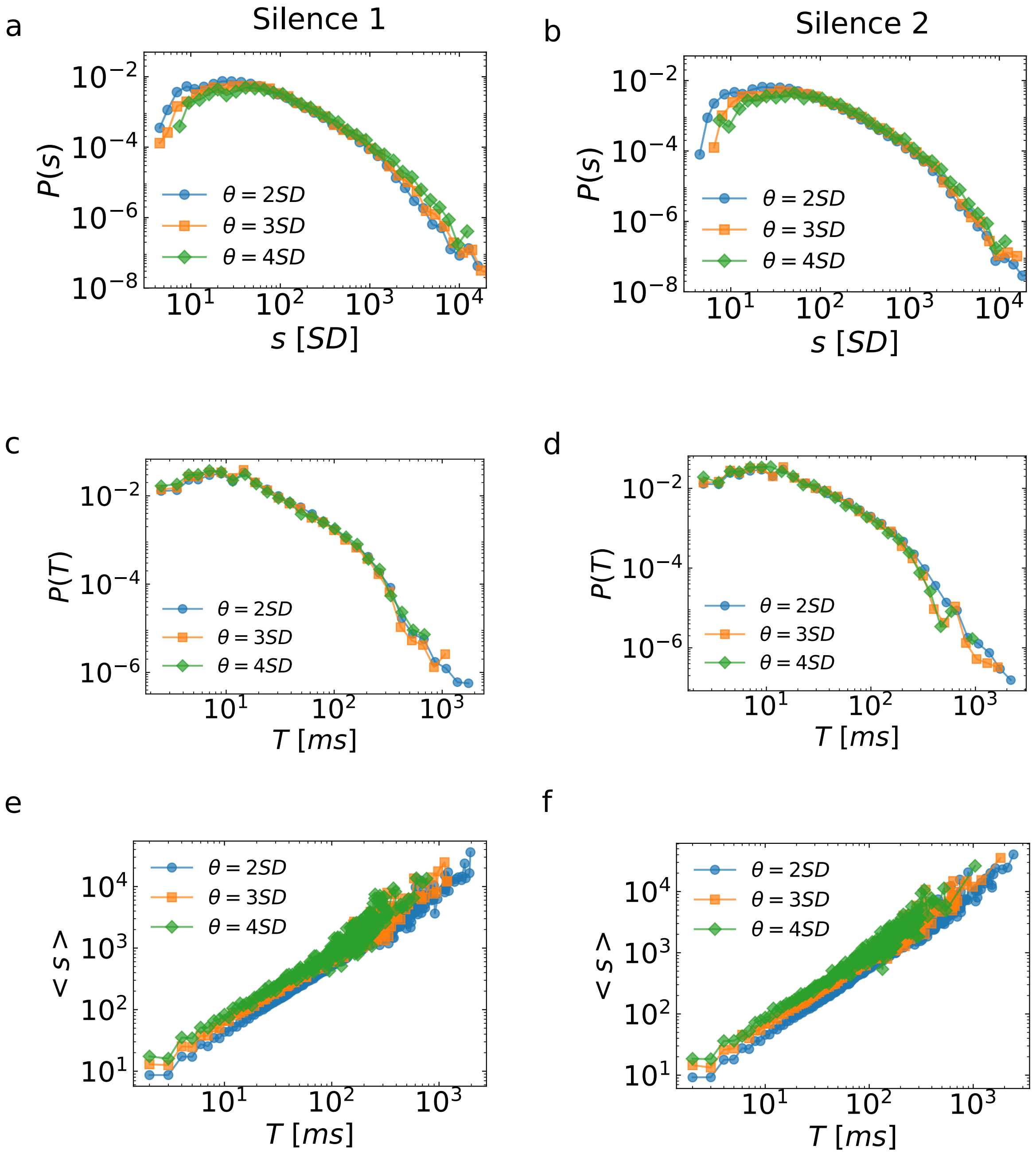


Fig. S8: **Avalanche dynamics is independent of the threshold $\theta$. a-b.** Avalanche size distribution, $P(s)$, in Silence 1 and Silence 2 (pooled data; $n = 33$). **c-d.** Avalanche duration distribution, $P(T)$, in Silence 1 and Silence 2 (pooled data; $n = 33$). **e-f.** Average avalanche size given a duration $\langle s \rangle \propto T^{\gamma}$ (pooled data; $n = 33$).

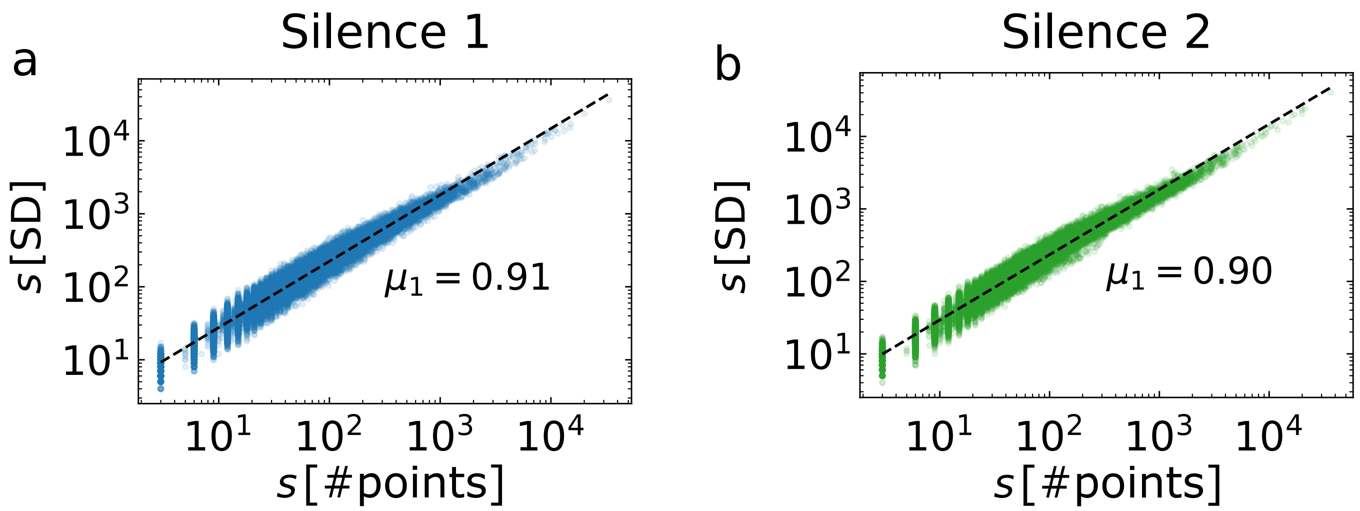


Fig. S9: **Relationship between the avalanche size $s$ and the number of events $n(t)\Delta t$.** Here we show the correspondence between the avalanche size $s(A_{cont})$ computed as the sum of the EEG amplitudes exceeding a threshold $\theta = 2SD$ as shown in Figure 1f and the avalanche size computed as the sum of the discrete events $s(A_{disc})$ exceeding a threshold $\theta = 2SD$ in a unit time $\Delta t = 2ms$ (the sampling time) as shown in Figure 3a. In both silences, we found the scaling relationship $s(A_{cont}) \propto (s(A_{disc}))^{\mu}$. **a.** Silence 1: $\mu_1 = 0.908 \pm 0.005$ (fit ± error of the fit). **b.** Silence 2: $\mu_2 = 0.900 \pm 0.001$.

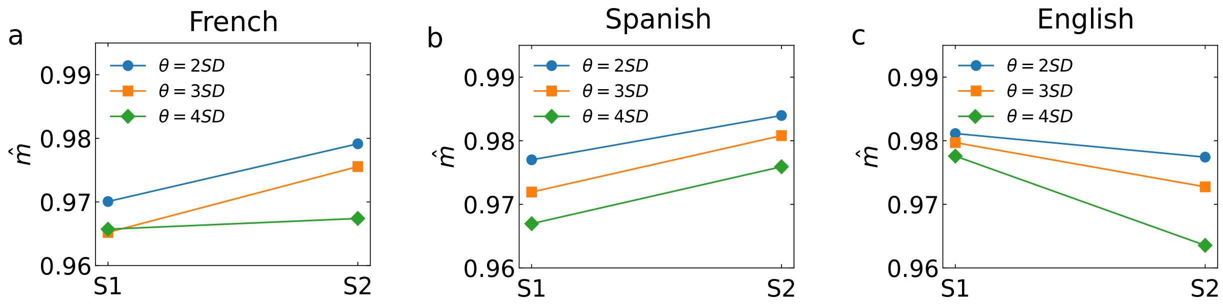


Fig. S10: **Branching parameter for different thresholds** $\theta$. **a.** French: $\theta = 2$ SD, $\hat{m}_1 = 0.970 \pm 0.005$ (mean ± SEM) and $\hat{m}_2 = 0.979 \pm 0.002$ ( pairwise $t$-test provides $p - value = 0.0307$); $\theta = 3$ SD: $\hat{m}_1 = 0.965 \pm 0.008$ and $\hat{m}_2 = 0.976 \pm 0.003$; $\theta = 4$ SD, $\hat{m}_1 = 0.966 \pm 0.009$ and $\hat{m}_2 = 0.967 \pm 0.005$. **b.** Spanish: $\theta = 2$ SD, $\hat{m}_1 = 0.977 \pm 0.002$ and $\hat{m}_2 = 9.984 \pm 0.001$ (pairwise $t$-test: $p = 0.0007$); $\theta = 3$ SD, $\hat{m}_1 = 0.972 \pm 0.002$ and $\hat{m}_2 = 0.981 \pm 0.003$ ( $p = 0.001$); $\theta = 4$ SD: $\hat{m}_1 = 0.967 \pm 0.002$ and $\hat{m}_2 = 0.976 \pm 0.005$. **c.** English: $\theta = 2$ SD, $\hat{m}_1 = 0.981 \pm 0.003$ and $\hat{m}_2 = 0.977 \pm 0.002$; $\theta = 3$ SD, $\hat{m}_1 = 0.980 \pm 0.003$ and $\hat{m}_2 = 0.973 \pm 0.003$ ($p = 0.019$); $\theta = 4$ SD, $\hat{m}_1 = 0.978 \pm 0.003$ and $\hat{m}_2 = 0.964 \pm 0.005$ ($p = 0.007$).

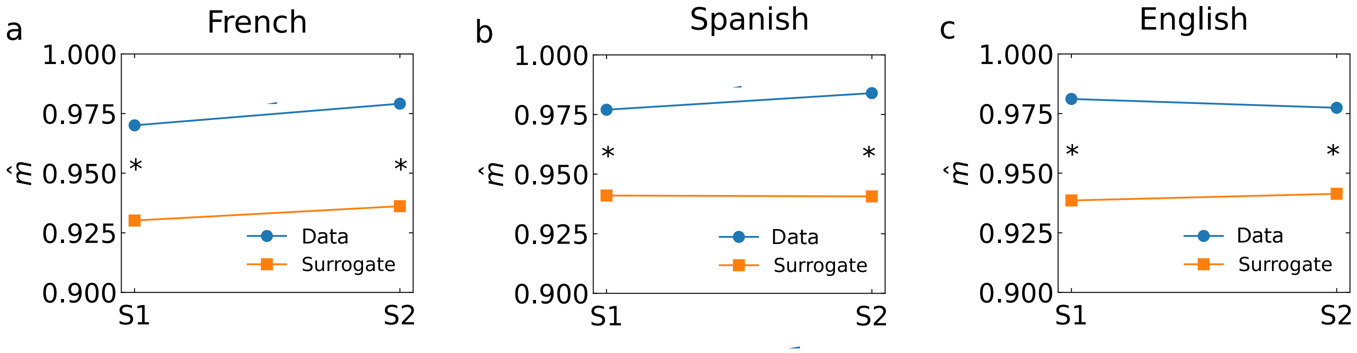


Fig. S11: **Branching parameter in surrogate data is significantly reduced.**. Average branching parameter calculated in surrogate data ($\hat{m}^{surr}$, orange) is compared with the values obtained for the original data ($\hat{m}$, blue) in Silence 1 (S1) and Silence 2 (S2). Black star indicates significant difference ($p < 0.05$) surrogate and data estimates. **a** French, Silence 1: $\hat{m}_1 = 0.970 \pm 0.005$ (mean ± SEM) and $\hat{m}_1^{surr} = 0.930 \pm 0.004$ ($p = 10^{-7}$). French, Silence 2: $\hat{m}_2 = 0.979 \pm 0.002$ and $\hat{m}_2^{surr} = 0.936 \pm 0.003$ (($p = 10^{-7}$)). No significative difference was found between $\hat{m}_1^{surr}$ and $\hat{m}_2^{surr}$. **b** Spanish, Silence 1: $\hat{m}_1^{nor} = 0.977 \pm 0.002$ vs $\hat{m}_1^{surr} = 0.941 \pm 0.003$ ($p = 10^{-7}$). Spanish, Silence 2: $\hat{m}_2^{nor} = 0.984 \pm 0.001$ vs $\hat{m}_2^{surr} = 0.941 \pm 0.001$ ($p = 10^{-7}$). No significant different was found between $\hat{m}_1^{surr} = 0.941 \pm 0.003$ and $\hat{m}_2^{surr} = 0.941 \pm 0.001$. **c** English, Silence 1: $\hat{m}_1 = 0.981 \pm 0.003$ vs. $\hat{m}_1^{surr} = 0.939 \pm 0.003$ ($p = 10^{-9}$). English, Silence 2: $\hat{m}_2 = 0.977 \pm 0.001$ vs. $\hat{m}_2^{surr} = 0.941 \pm 0.002$ ($p = 10^{-9}$).

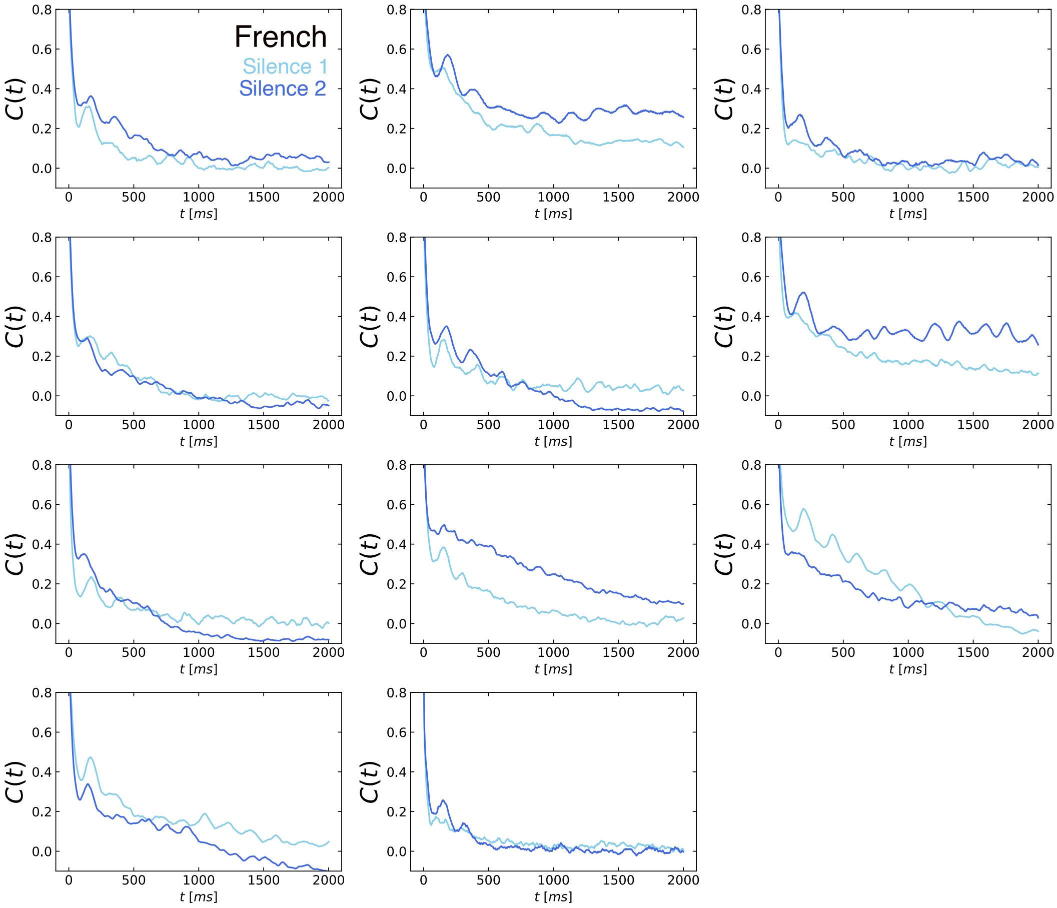


Fig. S12: **Autocorrelation of the instantaneous network activity in the French group during Silence 1 and Silence 2.**

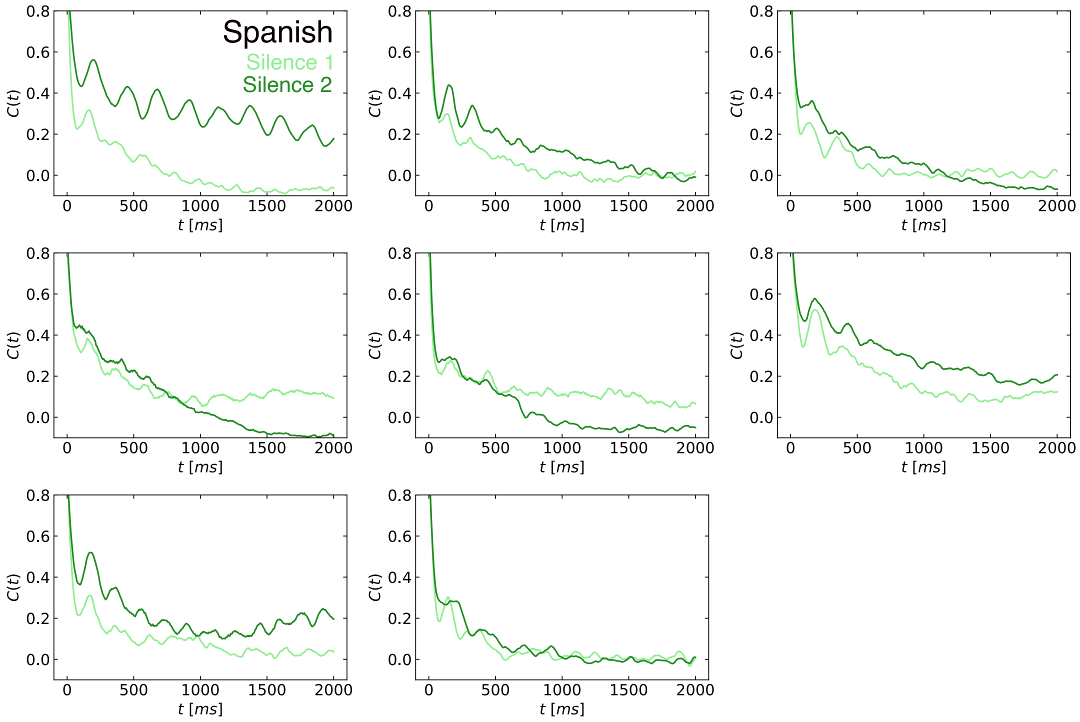


Fig. S13: **Autocorrelation of the instantaneous network activity in the Spanish group during Silence 1 and Silence 2.**

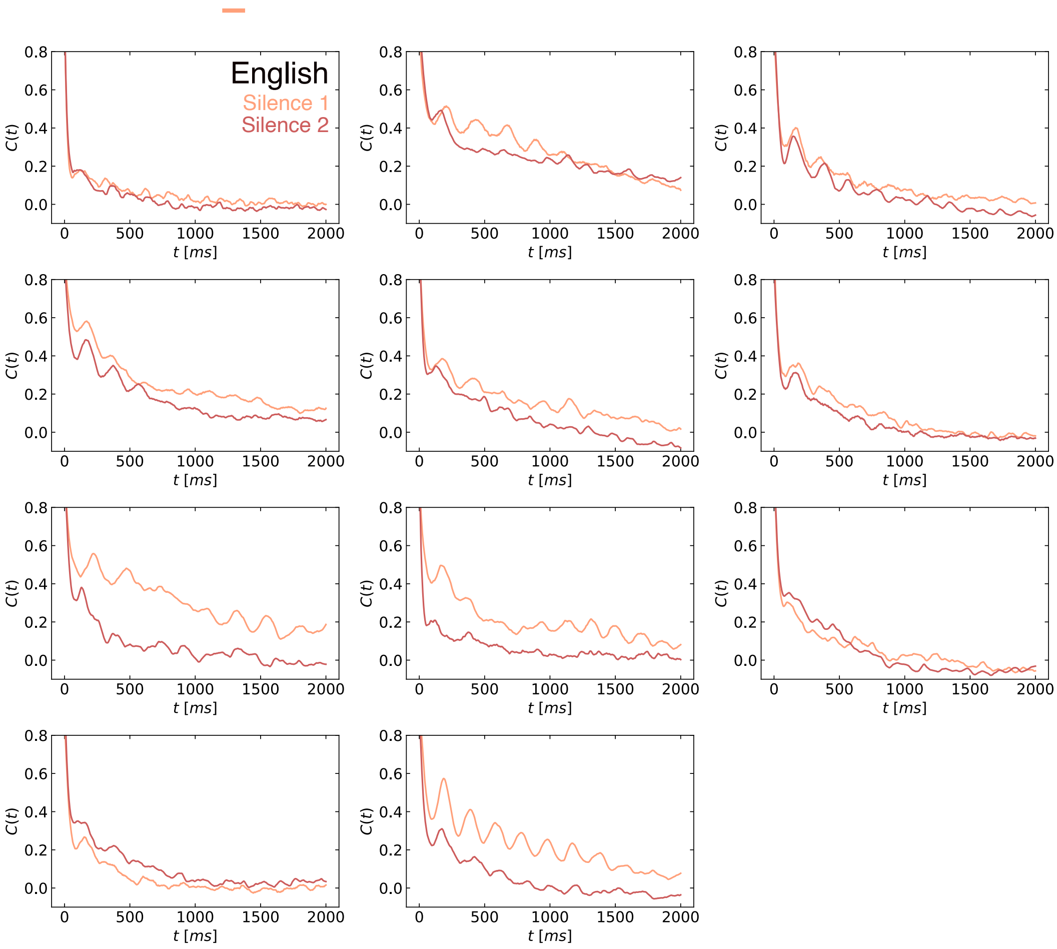


Fig. S14: **Autocorrelation of the instantaneous network activity in the English group during Silence 1 and Silence 2.** Unlike in the French and, to a lesser extent, in the Spanish group, the autocorrelation in Silence 2 is either weaker and decaying faster or unchanged compared to Silence 1.

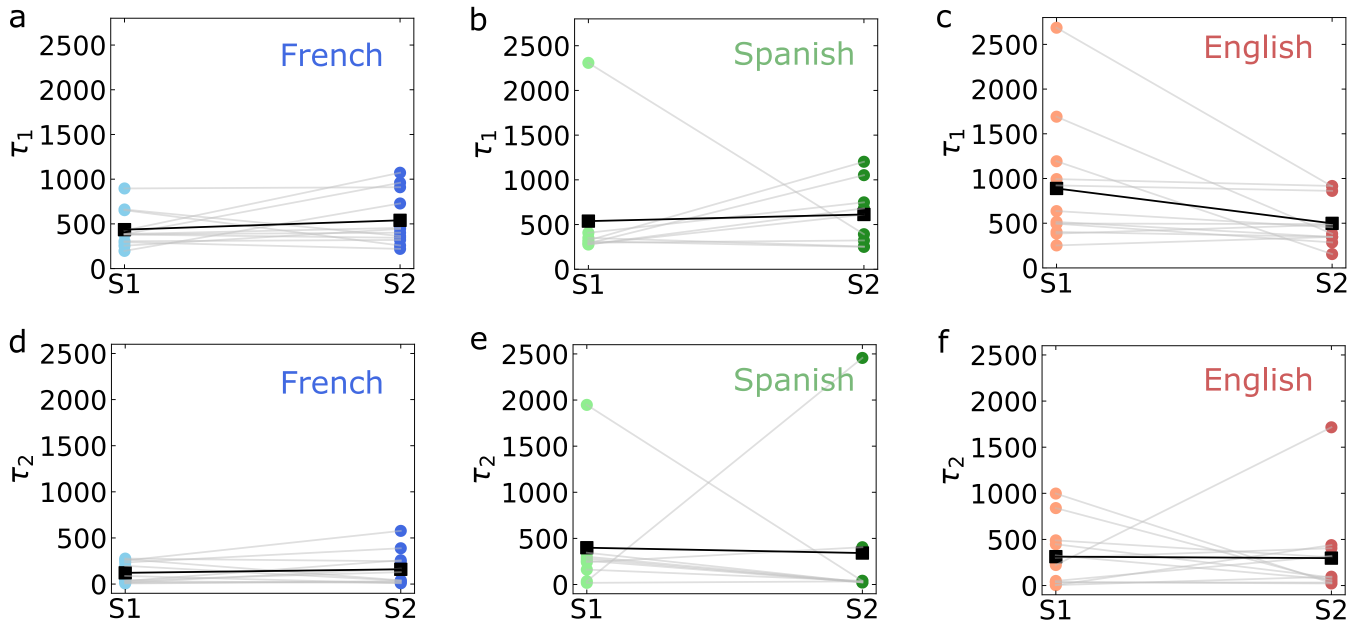


Fig. S15: **Time scales of the instantaneous network activity during Silence 1 and Silence 2.** We fit the autocorrelation of instantaneous network activity, $C(t)$, with the function $C_{fit}(t) = Ae^{-\frac{t}{\tau_1}} + Be^{-(\frac{t}{\tau_2})^{\gamma}} cos 2\pi \nu t + O$, where $\tau_1$ and $\tau_2$ are time constants and $\nu$ is the oscillation frequency. **a-c**. The time constant, $\tau_1$, of the exponential decay of the autocorrelation in Silence 1 and Silence 2 for the three language groups. In French and Spanish we notice that $\tau_1$ tends to increases in Silence 2 ($t$-test: French, $p = 0.29$; Spanish, $p = 0.79$). For English instead, $\tau_1$ decreases in Silence 2 (c. $t$-test: $p = 0.05$). **d-f**. Time constant, $\tau_2$, of the stretched exponential decay associated to long time scales of the autocorrelation in Silence 1 and Silence 2 for the three language groups.

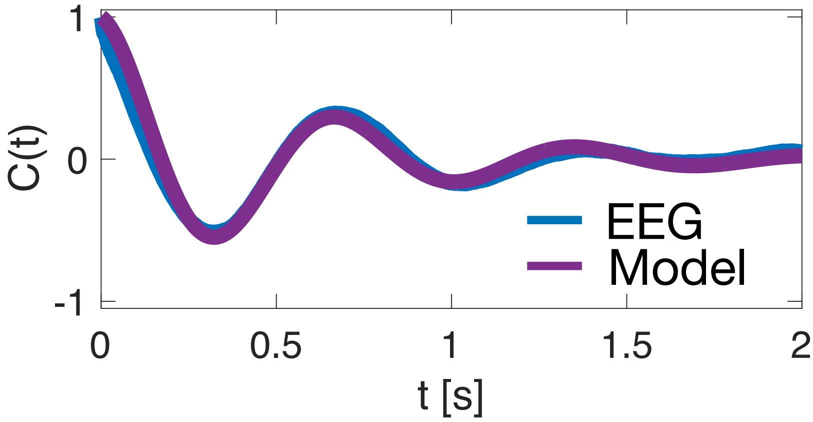


Fig. S16: **Example of a model fit for the empirical autocorrelation of an EEG signal.**

### Supplementary tables

| | French | | Spanish | | English | |
|---|---|---|---|---|---|---|
| | $\tau$ | $\alpha$ | $\tau$ | $\alpha$ | $\tau$ | $\alpha$ |
| R | 208.5 | 100.8 | 74.9 | 80.0 | 44.4 | 34.5 |
| p-value | $8 \times 10^{-16}$ | $5.0 \times 10^{-8}$ | $2.7 \times 10^{-6}$ | $5 \times 10^{-9}$ | 0.005 | $3.9 \times 10^{-4}$ |

Table 1: Likelihood ratio $R = lnL_p/L_e$ between power-law and exponential fit for the avalanche size and duration distributions in Silence 1.

| | French | | Spanish | | English | |
|---|---|---|---|---|---|---|
| | $\tau$ | $\alpha$ | $\tau$ | $\alpha$ | $\tau$ | $\alpha$ |
| R | 91.7 | 84.3 | 85.3 | 21.5 | 70.9 | 19.4 |
| p-value | $7.8 \times 10^{-7}$ | $4.4 \times 10^{-7}$ | $1.6 \times 10^{-6}$ | 0.011 | $7.4 \times 10^{-5}$ | 0.05 |

Table 2: Likelihood ratio $R = lnL_p/L_e$ between power-law and exponential fit for the avalanche size and duration distributions in Silence 2.

| | $\bar{\tau}$ | $\bar{\alpha}$ | $\bar{R}(\tau,\alpha) = \frac{\bar{\alpha}-1}{\bar{\tau}-1}$ | $\bar{\gamma}_{fit}$ | $\lvert\bar{R}(\tau,\alpha) - \bar{\gamma}_{fit}\rvert$ | $p - value$ |
|---|---|---|---|---|---|---|
| French S1 | 1.522 ± 0.043 | 1.965 ± 0.099 | 1.847 ± 0.244 | 1.164 ± 0.028 | 0.683 ± 0.328 | 0.025 |
| French S2 | 1.499 ± 0.054 | 1.612 ± 0.080 | 1.225 ± 0.208 | 1.210 ± 0.018 | 0.015 ± 0.190 | 0.430 |
| Spanish S1 | 1.425 ± 0.034 | 1.536 ± 0.075 | 1.263 ± 0.204 | 1.165 ± 0.026 | 0.098 ± 0.118 | 0.540 |
| Spanish S2 | 1.416 ± 0.041 | 1.742 ± 0.058 | 1.789 ± 0.227 | 1.212 ± 0.013 | 0.568 ± 0.133 | 0.002 |
| English S1 | 1.492 ± 0.055 | 1.781 ± 0.099 | 1.585 ± 0.262 | 1.161 ± 0.025 | 0.424 ± 0.114 | 0.004 |
| English S2 | 1.775 ± 0.324 | 1.370 ± 0.046 | 1.657 ± 0.086 | 1.210 ± 0.014 | 0.565 ± 0.354 | 0.040 |

Table 3: Average values of the critical exponents and scaling relationship for individual subjects in different language groups during Silence 1 and Silence 2. Errors represent SEM. The $p$-value on the last column refers to the difference between $\bar{R}(\tau,\alpha)$ and $\bar{\gamma}_{fit}$.
